# What canonical online and offline measures of statistical learning can and cannot tell us

**DOI:** 10.1101/2021.04.19.440449

**Authors:** Ava Kiai, Lucia Melloni

## Abstract

Statistical learning (SL) allows individuals to rapidly detect regularities in the sensory environment. We replicated previous findings showing that adult participants become sensitive to the implicit structure in a continuous speech stream of repeating tri-syllabic pseudowords within minutes, as measured by standard tests in the SL literature: a target detection task and a 2AFC word recognition task. Consistent with previous findings, we found only a weak correlation between these two measures of learning, leading us to question whether there is overlap between the information captured by these two tasks. Representational similarity analysis on reaction times measured during the target detection task revealed that reaction time data reflect sensitivity to transitional probability, triplet position, word grouping, and duplet pairings of syllables. However, individual performance on the word recognition task was not predicted by similarity measures derived for any of these four features. We conclude that online detection tasks provide richer and multi-faceted information about the SL process, as compared with 2AFC recognition tasks, and may be preferable for gaining insight into the dynamic aspects of SL.

## Introduction

Statistical learning (SL) refers to the ability to extract statistical regularities from the sensory environment. SL is thought to be critical for the segmentation of continuous sensory information, and the discrimination and prediction of stimulus items. [1],[2] As such, it is considered a key mechanism in language acquisition, [3] and other high-level aspects of cognitive and motor learning. [4] In the language domain, SL is widely believed to play a pivotal role in learning to distinguish the boundaries of words in a continuous speech stream, by leveraging information about the transitional probabilities of individual syllables through repeated exposure. [5],[6]

Since SL is cumulative in nature, a major challenge in the field has been to develop appropriate measures to quantify the time course of learning, as well as to determine which precise features of the continuous speech stream have been learned. For example, whether regularities have been extracted merely at the level of transitional probabilities, the ordinal position of the syllables within word units, or whole words (“chunks”). [7]

Current tools assessing learning rely primarily on explicit behavioral testing administered after the learning session. Typically, these “offline” tasks include a forced-choice task in which participants must discern which of two alternatives is the most familiar sequence of stimulus items: a sequence that is consistent with the learned regularities (e.g. a “pseudoword” in the artificial language of the speech stream) and a foil which violates the learned regularities, but consists of the same stimulus items (e.g. a non-word). SL has been demonstrated in both adults and infants in the auditory [8]–[12] and visual [13],[14] modalities. Although many studies report that participants can successfully discriminate lawful sequences in these tasks after only a few minutes of exposure, performance is typically just barely above chance (~60%). [5],[10],[15],[16] In addition, some argue that the explicit testing by itself could introduce unwanted interference effects, as participants’ memory of the lawful structures is vulnerable to distortion through repeated exposure to foil sequences. Consequently, explicit offline tasks are often limited to 16 or 32 trials, and may suffer from confounding and lack of statistical power. [17]

More recently, studies have examined SL through a combination of tasks, including some from of “online” task. These online tasks have the potential to provide a more sensitive measure of SL, important for understanding the dynamic aspects of extracting regularities from continuous input (e.g. how early SL occurs, how robust or stable it is over the course of the exposure phase, and whether it varies by stimulus). Typically, online SL tasks (“target detection tasks”) entail (1) asking participants to detect a target stimulus via keypress while being presented with a continuous stimulus stream, and (2) measuring reaction time (RT) to targets. Studies employing this task in both visual [13],[18],[19] and auditory [17],[20],[21] domains report that RTs to targets are modulated by the predictability of the target, suggesting that sensitivity to statistical regularities facilitates the speed of detection. Targets with lower transitional probabilities elicit longer RTs than targets with higher transitional probabilities. This RT effect can be observed as early as the second presentation of a target stimulus. [15] Similarly, in Gómez et al., participants detected click sounds embedded in continuous speech more rapidly when clicks occurred between rather than within pseudowords, suggesting that the stronger predictions generated by the learned pseudoword units interfered with click detection and thereby incurred longer RTs [22] (but see [23], for a non-replication). Though implicit online tasks tend not to suffer from the shortcomings of offline tasks mentioned in the previous paragraph, they can be more difficult to interpret, as distinct features of the stimulus stream (e.g. co-occurrence frequency and transitional probability) co-vary in many study designs but may differ in their contribution to the observed results. New evidence has called into question the validity of some previous findings that used RTs to measure SL. [13],[24] Specifically, Himberger et al. have demonstrated that RT facilitation could be observed independently of any regularity-learning if the task design confounds the position of the target within a triplet sequence with its position in the test stream. [25] The authors argued that RTs to targets in the first position of a triplet may be slower than those to targets in later positions due to trivial facilitation, as participants became more familiar with the task.

While there is ongoing debate as to whether SL is a unitary or multi-dimensional process, involving successive computational steps, numerous studies have tested participants on several learning tasks within the same experiment, to assess whether performance correlates among diverse measures. These studies have yielded conflicting empirical evidence as to whether detection-type online SL tasks can predict performance in the canonical 2AFC task. Earlier studies have reported significant, but small correlations (e.g. *ρ* = 0.42 [21], ρ = 0.46 [26]) between online (target detection or analogous tasks) and offline measures (2AFC discrimination or recognition tasks). Meanwhile, a larger number of studies report no correlation between these measures. [11],[18],[20],[23],[25]

The weak to nonexistent correlation between these SL measures has been largely discussed on a theoretical level. [18],[22] (But see [20] for an empirical treatment of the subject.) Some authors note that the weak relationship could be due to the simple fact that the two tasks rely on distinct cognitive processes, as they purportedly engage a different types of memory (implicit vs. explicit). Specifically, the target detection tasks are thought to tap onto implicit knowledge of the regularities, while the recognition tasks demand that information gleaned from the stream be made explicit. [20]. Alternatively, the weak relationship may also be ascribed to the different psychometric sensitivity of the two tasks: target detection tasks typically test all stimulus items over a longer test period, resulting in a comparatively larger number of trials. Meanwhile, word recognition tasks are often designed to test memory for the higher-level units (e.g. tri-syllabic pseudowords), and rarely exceed 36 trials in total. [12]

In this study, we addressed the question of why these two measures might be uncorrelated or weakly correlated, despite strong evidence that both tasks are indeed sensitive to the learning of embedded regularities. We hypothesized that the online and offline tasks might tap onto distinct features of the structured sequences presented during exposure, and that differences in task sensitivity to these features might explain the weak to non-existent correlation between the two tasks.

We tested participants in an online target detection task during exposure to a continuous stream of speech syllables, followed by a standard 2AFC pseudoword vs. part-word recognition task, and compared individual performance across both tasks (Experiment 1). We then aimed to replicate the results of the target detection task, and additionally included a control condition in which participants performed the detection task during exposure to streams of randomly ordered syllables (Experiment 2). Finally, to better understand which features of the syllable stream (transitional probability, triplet position, word grouping, or duplet pairing) are captured by RT data, we performed a representational similarity analysis (RSA) on group-level and participant level data. By providing rich information about what information is contained in RTs, RSA is a powerful tool that can provide novel insight into the empirical disparity between implicit online and explicit offline tests of SL. Our results suggest RT data from the target detection task reflects learning of several sequence features, but similarity measures for these features fail to predict word recognition performance.

## Results

### Experiment 1

In Experiment 1, participants were exposed to streams of continuous speech syllables (consisting of four repeating tri-syllabic pseudowords, made up from a bank of 12 unique syllables). During exposure, participants were asked to detect via keypress a target syllable in the auditory stream. Before each of 24 (~1 minute long) trials they heard a different target syllable. After the exposure phase, participants performed a 2AFC word recognition task in which they heard each of the four pseudowords presented alongside four part-words, and reported which of each test pair they believed was a word in the “alien language” they just heard.

Detection accuracy in the online target detection task (*M* = 0.70, *SD* = 0.46, *t*(32) = 10.19, *p* < 0.001), and the true positive rate ((*True Positives*)/(*True Positives + False Positives*) = 0.87) were high, indicating that participants (N = 33) paid attention to the stream and followed instructions. (See Supplementary Materials, **Fig. S1**, and **Table S3** for further analysis of detection accuracy as a function of triplet position, pseudoword, and target syllable.)

#### Triplet Position Modulates Reaction Time

To investigate whether subjects exhibited sensitivity to the statistical regularities in the stream, we evaluated the effect of target syllables’ position (word-initial, word-medial, or word-final) within a pseudoword on RT. To this end, we used a generalized linear mixed effects model (GLMM) with triplet position as predictor and RT (in seconds) as outcome variable. We found that RTs were modulated by the triplet position of the target syllable within the pseudoword (*X*^2^(2, *N* = 33) = 523.49, *p* < 0.0001, Type II Wald Chi-square test (hereafter “Type II”)). (Fig. 1a-b). To explore the differences in RT across the three triplet positions, we performed post-hoc pairwise comparisons on estimated marginal means for each position, with Tukey adjustment. RTs to word-initial syllables (position 1) (*M* = 524 *ms*, *SD* = 179 *ms*) were notably slower than those to word-medial (position 2) (*M* = 452 *ms*, *SD* = 164 *ms*; *z* = 15, *p* < 0.0001, *Cohen’s d* = 0.46) and word-final syllables (position 3) (*M* = 423 *ms*, *SD* = 161 *ms*, *z* = 22.72, *p* < 0.0001, *d* = 0.67). The difference in mean RT between word-medial and word-final positions was small but still significant (*z* = 7.46, *p* < 0.0001, *d* = 0.2).

**Figure 1. Exp. 1:**
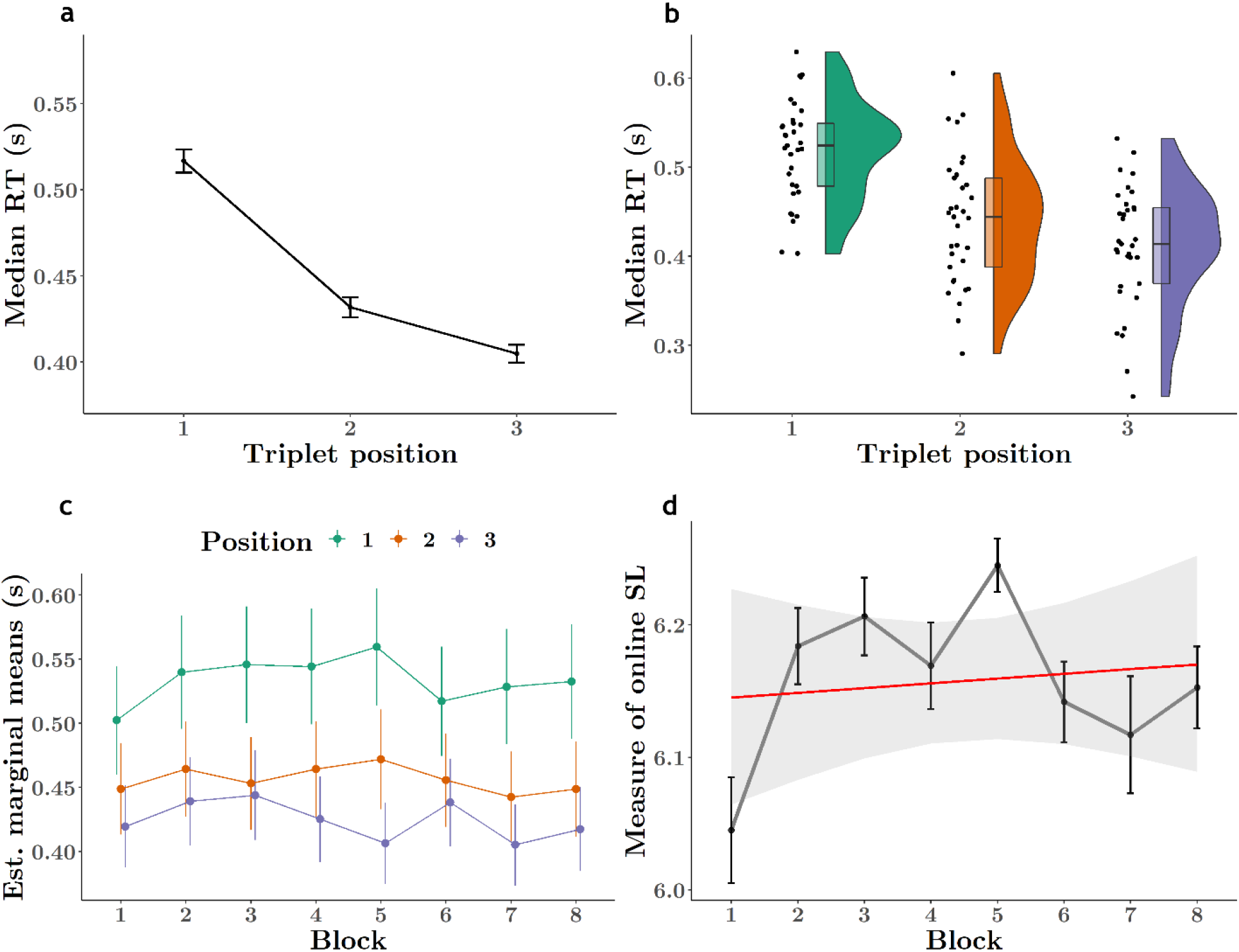
Online target detection reveals rapid and robust sensitivity to embedded regularities. A. Median reaction times (RT) to target syllables are modulated by the syllable’s triplet position in pseudowords. Participants responded more slowly to syllables in the word-initial (1^st^) position than to syllables in the word-medial (2^nd^) or word-final (3^rd^) position. Error bars represent 95% confidence intervals. B. Distribution of median RTs to each triplet position for each participant (black dots). (Jittered along x-axis for visibility.) Box plots indicate group median and 95% CI. C. RTs to targets in the 1^st^ vs 2^nd^ or 3^rd^ position are somewhat distinct in the first block (first 3 minutes of exposure), and clearly differentiated thereafter. Points represents estimated marginal means from the GLMM, vertical lines represent 95% CI. D. The magnitude of the SL effect (log mean RT to 1^st^ position – log mean RT to (2^nd^ & 3^rd^) position syllables) was smallest in the first block, but rose dramatically in the second block, thereafter hovering around the overall mean. (No difference in mean RTs to the three positions would result in a value of 0.) We observed no significant effect of block on this metric, suggesting that the modulation of RT as a function of target syllable position in the pseudowords had already occurred in the first block.

#### Rapid Onset of Graded RT to Predictable Syllables

Previous studies have found that a few minutes of exposure [27] or even a few occurrences of an embedded target syllable [15] were sufficient for participants to pick up the statistical structure, as measured by offline word recognition and RT, respectively, while longer exposures have been shown to provide little added benefit in the former task. [10] To relate to those findings we directly tested whether the graded RT effect emerged over the course of several blocks, or was present from the first block. We computed an ANOVA on our fuller model (see **Table S1**) with factors block (8) and triplet position (3) as predictors and RT (in seconds) as outcome variable. We found a significant interaction between block and triplet position (*X*^2^(14, *N* = 33) = 28, *p* = 0.014, Type II). Previous studies have revealed differentiation of responses in the online SL task as early as the second presentation of the pseudoword, [15], as well as above-chance offline word recognition after only five minutes of exposure in adults. [10] Since each of our blocks are ~ 3 minutes long, we predicted on the basis of these prior results that participants would have extracted the embedded regularities of the stream within the first two blocks, and focused our follow-up analyses on the first two blocks only.

To specifically examine the change in the RT pattern between the first two blocks, we performed an ANOVA on a model identical in structure with that above, but using only data from blocks 1 and 2. Here, we observed no interaction, but significant main effects for both block (*X*^2^(1, *N* = 33) = 8.95, *p* = 0.003, Type II) and triplet position (*X*^2^(2, *N* = 33) = 92.67, *p* < 0.0001, Type II) factors. The effect of block was driven by a small, overall increase in RT between blocks 1 and 2 (*M*_*block 1*_ = 443 *ms*, *SD* = 159 *ms*; *M*_*block 2*_ = 476 *ms*, *SD* = 170 *ms*; *z* = −3.09, *p* = 0.002, *d* = −0.07), while contrasts between each position revealed a graded RT effect that was largest for position 1-2 (*M*_*position 1*_ = 506 *ms*, *SD* = 175 *ms*; *M*_*position 2*_ = 458 *ms*, *SD* = 157 *ms*; *z* = 6.24, *p* < 0.001, *d* = 0.19) and 1-3 mean differences (*M*_*position 3*_ = 429 *ms*, *SD* = 157 *ms*; *z* = 9.01, *p* < 0.001, *d* = 0.27), but also significant for position 2-3 (*z* = 3.1, *p* = 0.006, *d* = 0.08). Since we did not find evidence that the RT effect was modulated by block (no interaction), we focused our next analysis on block 1 only. An ANOVA performed on a model using data only from block 1 and therefore only triplet position as predictor revealed again a main effect of triplet position (*X*^2^(2, *N* = 33) = 27.93, *p* < 0.0001, Type II) and, between each level, the same graded RT effect (*position* 1 − 2: *z* = 2.89, *p* = 0.01, *d* = 0.12; *position* 2 − 3: *z* = 2.45, *p* = 0.039, *d* = 0.21; *position* 1 − 3: *z* = 5.09, *p* < 0.0001, *d* = 0.08). This corroborates our previous finding that the factor block did not contribute significantly to model fit (*X*^2^(1, *N* = 33) = 2.12, *p* = 0.15): reaction times differentiate early on and persist across the whole data set. (**Fig. 1c**)

To relate to previous studies [26], we computed a measure of online statistical learning, borrowed from Siegelman and colleagues, as follows:

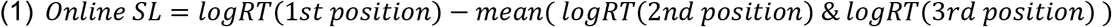

A linear model with the SL measure as outcome variable and block as predictor failed to reveal any effect of block on this composite measure (*X*^2^(7, *N* = 33) = 6.50, *p* = 0.48, Type II). (**Fig. 1d**) This is an additional piece of evidence in support of early emergence of differentiated reaction times and stable behavior thereafter.

Lastly, we wanted to address a possible confound that could have helped generate the RT effect. As argued by Himberger et al. [25], the widely observed graded RT effect could be driven by a trivial, overall speeding-up of RTs combined with the fact that targets appear later in the stream as well as later in the pseudoword or triplet. To ensure that this confound is not present in our data, we ran a linear model with triplet position and stream position (1-216) within each trial as predictors. While we observed a main effect of triplet position (*F*(2) = 362.44, *p* < 0.001), we observed no effect of stream position (*F*(1) = 0.0029, *p* = 0.96). (See Supplementary Materials and **Fig. S2**.) We also tested to see if RTs decrease monotonically over single trials. A linear model with block and target number (number of occurrences of each target within each trial) revealed no interaction of the factors on RT (in seconds), but main effects of each. Given that we already addressed the effect of block, we re-ran the model with only target number as predictor, which again revealed a main effect (*F*(17) = 3.2, *p* < 0.0001). However, we observed that the variation in RT between target numbers was not unidirectional, but wavered above and below the mean RT. (**Fig. S2**)

#### Pseudowords Can Be Distinguished From Part-words

After the exposure phase, participants performed a 2AFC word recognition task in which they heard each of the four pseudowords presented alongside four part-words, and reported which of each test pair they believed was a word in the alien language they just heard. Participants correctly distinguished the pseudowords from the part-word foils significantly above chance (chance level = 50%, or 8 out of 16 trials) (*M* = 0.62, *SD* = 0.2; *t*(37) = 3.78, *p* < .001, *d* = 0.61), indicating that participants were sensitive to the implicit regularities of the syllable stream and able to use this information to explicitly discriminate pseudowords from sequences of syllables that crossed word boundaries. **(Fig. 2a)** 71% of participants (27 out of 38) completed the task with a mean accuracy greater than chance (50%, 8 out of 16 trials). To rule out the possibility that above-chance word recognition was driven by any particular word, and thus potentially an artifact of the stimulus materials, we performed an exploratory analysis in which we calculated the proportion of correct responses for each pseudoword individually (out of 4 trials). We found that across participants, 3 out of the 4 pseudowords were discriminated above chance (2 out of 4 trials), indicating that the above-chance performance can be attributed to learning of the underlying structure (*t*_*mipola*_ (37) = 3.24, *p* = 0.01, *t*_*nugadi*_ (37) = 3.31, *p* = 0.008, *t*_*rokise*_ (37) = 0.36, *p* = 1.0, *t*_*zabetu*_ (37) = 2.99, *p* = 0.02, Bonferroni corrected for four comparisons). (**Fig. S3**)

**Figure 2. Exp. 1:**
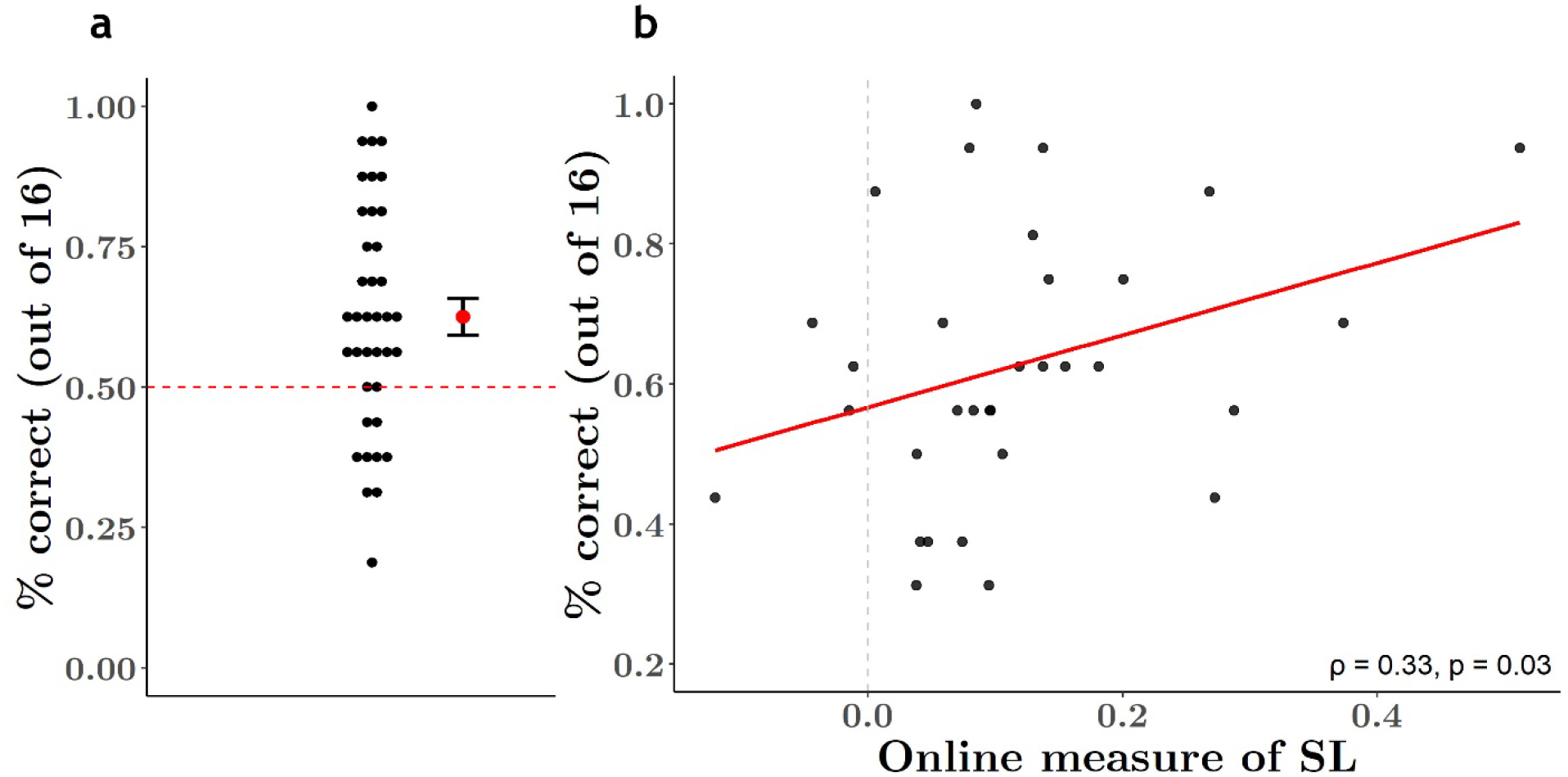
Participants discriminated pseudowords from part-words, but online and offline SL was only weakly correlated. *A.* Word recognition performance was above *the 0.5* chance level *(dashed red line)*, suggesting participants were able to use implicitly learned regularities during the exposure phase to explicitly discriminate pseudowords from part-word foils. *Red dot and* error bars represent *group* mean *(0.62)* and *SEM (0.2)*. *Black dots represent percent correct trials (out of 16) for each individual. B. Pearson correlation of each individual’s word recognition performance (percent correct trials) and RT effect in the target detection task (online measure of SL). Performance in the two tasks was weakly correlated (ρ* = 0.33, *p* = 0.03*). Dashed grey line represents threshold at which there was no difference in mean RT to targets in the three positions: those with values above 0 showed faster responses to 2*^nd^ *and 3*^rd^ *position targets vs. 1*^st^ *position targets, those with values below 0 showed the opposite effect.*

#### Online and Offline Measures Are Weakly Correlated

Next, we calculated the measure of online SL (see equation 1 above) for each participant and correlated each individual’s online SL scores with their word recognition accuracy. Since we did not have complete data for all participants (see Methods), for this analysis we used data only from those participants with complete data in both tasks (N = 32). We found a weak but significant correlation between these two values at the 5% alpha level (*ρ* = 0.33, *t*(30) = 1.93, *p* = 0.03, Pearson’s product-moment correlation, one-sided). (**Fig. 2b**)

To relate this finding to previous literature, we repeated this analysis using the procedure from [21] to compute the “RT score”: *median* (*RT*_1*st position*_) − *median* (*RT*_3*rd position*_). This analysis revealed an even weaker correlation, which did not reach statistical significance (*ρ* = 0.23, *t*(30) = 1.3, *p* = 0.1, Pearson’s product-moment correlation, one-sided). (**Fig. S4**) Finally, to see if certain syllable pairs might better predict word recognition accuracy, we considered the correlation between word recognition accuracy and the median difference between each triplet position pairing (i.e. 1-2, 2-3, and 1-3) in milliseconds. To obtain “RT scores” that are comparable between participants, we z-normalized RT values for each participant, computed median RTs to each triplet position, and computed the difference between the scaled median RTs for each position pairing for each participant. These values were correlated against the participant’s word recognition accuracy. We again found insignificant correlations between the two measures for all pairs (1 − 2: *ρ* = 0.22, *t*(30) = 1.24, *p* = 0.1; 2 − 3: *ρ* = 0.07, *t*(30) = 0.43, *p* = 0.33; 1 − 3: *ρ* = 0.23, *t*(30) = 1.3, *p* = 0.1, Pearson’s product-moment correlation, one-sided). (**Fig. S4**)

#### Discussion

Our study replicated two tasks that measure SL in distinct ways. Our offline word recognition task revealed a well-established effect of learning, where pseudowords marked by dips in transitional probability are discriminated from foil sequences of syllables that span across word. Likewise, our online target detection task revealed results consistent with previous literature: targets in predictable locations (i.e. word-medial and word-final positions, transitional probability = 1) elicited faster RTs than targets in less predictable locations (word-initial positions, transitional probability = 0.33). The rapid onset of this graded RT effect corroborates previous findings showing SL to be a fast and robust mechanism. Indeed, RTs differences by triplet position emerged during the first block, and remained relatively stable throughout the remainder of the experiment. Finally, this graded RT effect was not confounded by a trivial speeding-up of RT or the position of targets in the syllable sequence.

Intriguingly, a tracking of transitional probability alone cannot account for the results. Indeed, if participants were sensitive only to transitional probability, we would expect to find a significant difference in RT to syllables in word-initial positions versus word-medial and word-final positions, but no difference between word-medial and word-final syllables, since the latter two have the same transitional probability (1). Rather, we find that RTs to word-final syllables are also significantly faster than RTs to word-medial syllables, suggesting that triplet position, duplet pairing, and/or an awareness of the pseudoword grouping provided an addition source of predictability to prepare the response to the final syllable.

We found that the two measures of SL were weakly correlated. This correlation reached significance in only one of the three methods we used to evaluate the relationship, despite the fact that we followed two methods from earlier papers that identified a significant relationship. [21],[26] The weak and variable correlation we observed, in combination with previous findings reporting variable to null results [11],[18],[20],[23],[25], calls into question whether these two measures of SL are in fact evaluating learning of the same or different features of the input.

Each of the syllables in the stream can be characterized by its membership to a specific group for each of the following features: transitional probability (1 or 0.33), triplet position (word-initial/position 1, word-medial/position 2, or word-final/position 3), word grouping (*nugadi, rokise*, *mipola,* or *zabetu*), and within-word duplet pairings (*nu-ga, ga-di, ro-ki, ki-se*, etc.). [7] Meanwhile, there is some ambiguity about which of these features is exploited to achieve each task. Indeed, both the RT effect (between word-initial and –medial syllables) and above-chance word recognition could be achieved by leveraging transitional probabilities alone. However, If these two tasks can be accomplished with distinct sources of information, and if RTs do not capture any information related to word or duplet pairing, this might explain the lack of correlation.

To address this issue, we collected data in a second experiment in which participants performed only the online detection task. We aimed to replicate the graded RT effect found in the previous study, and to validate the online detection task as a genuine measure of learning. Experiment 2 consisted of an exposure phase/target detection task with a random stream consisting of the same stimulus materials, but no statistical regularities. This manipulation would allow us to confirm that our reported effects were driven primarily by the statistical regularities in the stream and not by unwanted variation in the stimuli acoustics. We then combined data from Experiments 1 and 2 to investigate feature coding in RTs though RSA; specifically, coding of transitional probability, triplet position, word grouping, and within-word duplet pairings.

### Experiment 2

As in Experiment 1, participants performed a target detection task while being exposed to a continuous speech stream. Before each of 12 (~1 minute long) trials, participants heard a target syllable, which they had to detect via keypress each time it appeared in the following stream. Condition order (structure, random) was counter-balanced across subjects, such that one-half performed the target detection task first for the structured stream and then for the random stream, and one-half performed the task in the opposite order. Each of the 12 syllables was tested once in each condition.

We first examined mean detection accuracy to ensure participants (N = 20) were engaged in the task. Overall detection accuracy (*M* = 0.82, *sd* = 0.38; *t*(19) = 13.48, *p* < 0.0001) and the true positive rate (*M* = 0.92) was high, confirming that participants adequately performed the task. Detection accuracy was numerically higher in the structured condition (*M* = 0.84, *sd* = 0.36) than in the random condition (*M* = 0.80, *sd* = 0.4), but this difference was not significant when tested through a one-sided test predicting the mean for the structured condition to be greater than random (*t*(36.2) = −1.37, *p* = 0.089). (See Supplementary Materials, **Fig. S1**, and **Table S4** for further analysis of detection accuracy as a function of triplet position, pseudoword, and target syllable.)

#### Triplet Position in Structured Stream Modulates Reaction Time

Using a GLMM with condition and target position as predictors, we observed an interaction between triplet position and condition (*X*^2^(2, *N* = 20) = 59.16, *p* < 0.0001, Type II). (See **Table S2** for regression results.) We then performed two planned contrasts. First, we evaluated the effect of triplet position within each level of condition to determine the modulation of RTs within each condition. As in Experiment 1, in the structured condition, we observed slower RTs to word-initial syllables (*M* = 580 *ms*, *SD* = 140 *ms*) than to word-medial syllables (*M* = 509 *ms*, *SD* = 133 *ms*; *z* = 12.69, *p* < 0.001, *d* = 0.56), and word-final syllables (*M* = 489 *ms*, *SD* = 128 *ms*; *z* = 16.37, *p* < 0.001, *d* = 0.73). The drop in mean RT between word-medial and word-final syllables was smaller but also significant (*z* = 12.42, *p* = 0.0002, *d* = 0.17). (**Fig. 3a-b**).

**Figure 3. Exp. 2:**
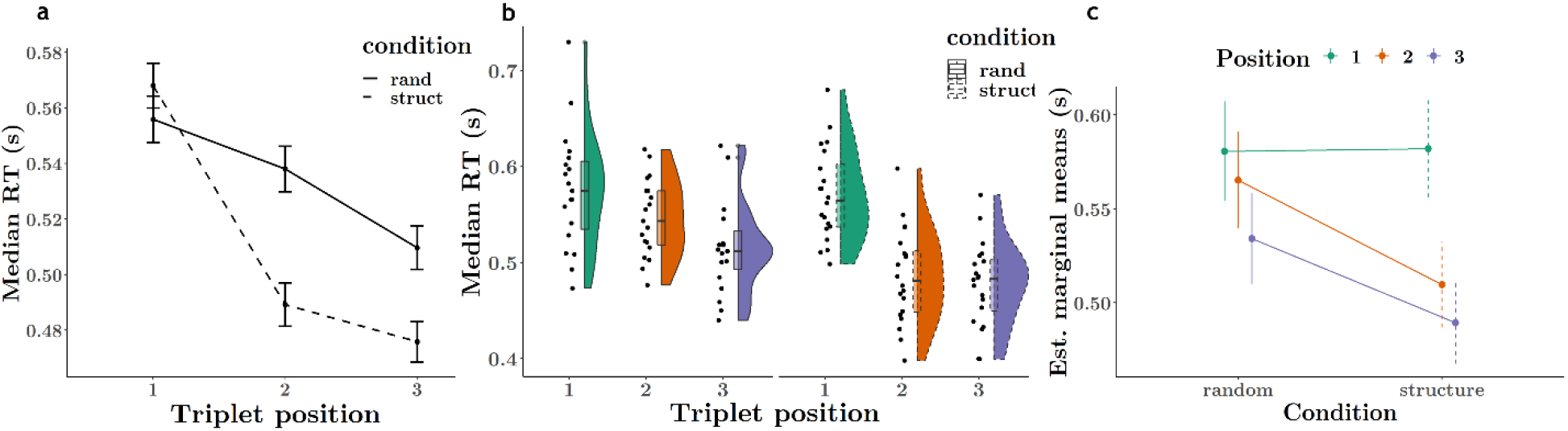
RT to target syllables modulated by triplet position and the presence of structure. A. In the structured condition (dashed line), the triplet position of target syllables modulated RT, replicating Exp. 1. Surprisingly, a less pronounced RT modulation by position was also observed in the random condition (solid line). Error bars represent 95% confidence intervals. B. Distribution of median RTs to each triplet position for each participant (black dots) in the random condition (left, solid violin & boxplot outline) and structured condition (right, dashed violin & boxplot plot outline). (Jittered along x-axis for visibility.) Box plots indicate group median and 95% CI. C. Estimated marginal means from GLMM with triplet position and condition as predictors for RT. RTs to target syllables in the 1^st^ position are equal between conditions, but RTs to those in 2^nd^ and 3^rd^ positions are significantly faster in the structured as compared with the random condition. This suggests the transitional probability structure was responsible for the RT effect, since less predictable syllables (1^st^ position, TP = 0.33) were responded to equally quickly, but predictable syllables (2^nd^ and 3^rd^ position, TP = 1), which appeared only in the structured stream, elicited markedly faster responses.

We observed a similar general RT pattern in the random condition, although with far smaller marginal differences. Here, mean RTs to word-initial (*M* = 579 *ms*, *SD* = 133 *ms*) were slower than those to word-medial (*M* = 563, *SD* = 132; *z* = 2.45, *p* = 0.04, *d* = 0.11) and to word-final (*M* = 534 *ms*, *SD* = 133; *z* = 7.86, *p* < 0.0001, *d* = 0.35) syllables as well. RTs to word-medial syllables were also slower than RTs to word-final syllables (*z* = 5.36, *p* < 0.0001, *d* = 0.24). (**Fig. 3a-b**) Given that there were no regularities in the random stream that could bias reaction times to certain tokens more than others, we hypothesized that the modulation observed here is due to slight variations in the discriminability of some CV syllables [28] and/or a weak carry-over effect of exposure to the structured stream before the random stream. (See Supplementary Materials, **Fig. S5**).

In addition, we ran a linear model using condition and target number (as a factor) as predictors to confirm that the potential confound of linearly decreasing RTs (in seconds) over the course of each trial/stream did not drive this effect. We found an interaction between condition and target number (*F*(17) = 2.39, *p* = 0.001). However, RTs in fact followed a quadratically-shaped pattern over the course of trials for both conditions, generally increasing until occurrence 12-14, and then decreasing. (**Fig. S2d**)

In our second contrast, we evaluated the effect of condition for each level of triplet position, i.e. how much condition affected RT to targets in each triplet position. We observed that the presence of structure significantly decreased mean RT for word-medial (*z* = 5.32, *p* < 0.0001, *d* = 0.43) and word-final targets (*z* = 4.54, *p* < 0.0001, *d* = 0.37). However, RTs to word-initial targets did not significantly vary between conditions. (*z* = −0.12, *p* = 0.90, *d* = −0.01) (**Fig. 3c**).

#### Discussion

In Experiment 2, we were able to replicate our main finding from Experiment 1 and establish that our observed RT effect reflects genuine learning of implicit structure. When a continuous speech stream featured implicit structure, RTs to predictable targets were significantly faster than those to less predictable targets. Importantly, RTs to word-initial syllables, which are unpredictable, were roughly equal between random and structured streams.

Himberger and colleagues recently argued that the graded RT effect observed in numerous SL studies is an artifact unrelated to the regularities that experimenters expect participants to learn, but rather a consequence of general RT facilitation, combined with a design that confounds position in the triplet with position in the stream. [25] We demonstrate empirically that this confound is not present in our data. RTs in both experiments did not trend towards linearly faster responses over the course of a trial/stream; RTs in Experiment 1 hovered around the mean, while RTs in both conditions in Experiment 2 increased for a majority of the trial. In addition, our design makes it unlikely that the RT effect would be vulnerable to confounding, as target syllables occurred across all 216 positions with each trial/stream in both experiments, making all targets equally susceptible to longer-term RT trends, Furthermore, while the authors’ critique may apply to certain uses of the online detection task [13],[24], other studies have employed the task while successfully controlling for any effect of stream position. [15]

### Feature Sensitivity in Online Target Detection

While we were able to show that exposure to structured streams resulted in a graded RT effect for target syllables in different triplet positions, the target detection task cannot directly tell us which feature of the target syllables (transitional probability, triplet position, word grouping, or duplet pairing) was “learned”, and responsible for the RT effect. To address this ambiguity, we performed a RSA on the RT data from the online target detection tasks to determine if the observed patterns of RTs could reveal sensitivity to any of the four specific features of the structured streams outlined here. RSA entails computing similarity matrices for responses to different stimuli, and allows one to observe which distinctions are emphasized in participant responses. [29],[30]

For this analysis, we combined the data from Experiments 1 (N = 33) and 2 (N = 20, structured condition only) for a total N of 53. Similarity matrices for each participant, obtained by correlating RTs to each syllable (12), were used to generate two subsets of the data for each of the four features outlined above. For each feature, we identified a *within* and an *across* group. *Within* groups consisted of the correlation values between all pairs of syllables characterized by that feature. *Across* groups consisted of correlations between pairs where the pairing violates the feature or represents the opposite feature type. (See **Fig. S6** for a graphical representation of the correlation values that entered into each group for each analysis. In the examples given below, letters (representing pseudowords) and numbered subscripts (representing triplet position) provide examples of the syllable pairs used in each analysis.) Finally, we assessed whether *within* or *across* group similarity was greater using a Wilcoxon rank sum test for each feature. A significant difference between *within* and *across* similarity values would suggest that the feature in question is represented in the RT data.

#### RTs Track Transitional Probability, Triplet Position, Word Groupings and Duplets

For the test of triplet position (N = 735), *within* values included the correlation between pairs of syllables with the same triplet position: all pairs word-initial syllables (e.g. A_1_-B_1_, B_1_-C_1_, etc.), all pairs of word-medial syllables (e.g. A_2_-B_2_, B_2_-C_2_, etc.), and all pairs of word-final syllables (e.g. A_3_-B_3_, B_3_-C_3_, etc.). *Across* values included correlations between all syllable pairs where each syllable of the pair had a different triplet position (e.g. A_1_-A_2_, A_1_-B_2_, C_2_-D_3_, etc.). Similarity *within* triplet positions was significantly lower than similarity *across*triplet positions (*Median difference* = −0.004, *V*(53) = 5334842911, *p* = 0.027, *Bonferroni corrected, CI* = [−0.007, −0.001], Wilcoxon rank sum test).

For the test of transitional probability (N = 250), *within* values included the correlation between all pairs of word-medial and word-final syllables, as all these syllables had a transitional probability of 1 in the exposure stream and appeared “within” pseudowords (e.g. A_2_-A_3_, A_2_-B_2_, A_2_-B_3_, etc.). *Across* values consisted of the correlation between all pairs of word-initial syllables, which had a transitional probability of 0.33 and appeared at pseudoword boundaries (e.g. A_1_-B_1_, B_1_-C_1_, etc.). Similarity was significantly higher for the *within* as compared with the *across* group. (*Median difference* = 0.07, *V*(53) = 721262654, *p* < 0.0001, *Bonferroni corrected, CI* = [0.07, 0.08], Wilcoxon rank sum test)

For the test of word grouping (N = 496), *within* values included the correlation between all pairs of syllables within each pseudoword (e.g. A_1_-A_2_, A_2_-A_3_, A_1_-A_3_, etc.). *Across* values included “phantom” word pairs where each syllable in the pair is drawn from different pseudowords (e.g. A_1_-B_2_, B_2_-C_3_, C_1_-D_2_, etc.). Similarity values were significantly higher *within* versus *across* word groupings. (*Median difference* = 0.015, *V*(53) = 2516837351, *Bonferroni corrected, CI*, *p* < 0.0001, *CI* = [0.011, 0.019], Wilcoxon rank sum test).

Finally, for the test of duplet pairing (N = 329), *within* values were correlations between all pairs of consecutive syllables within pseudowords (e.g. A_1_-A_2_, A_2_-A_3_, etc.), while *across* values were correlations between all other syllable pairs that were heard adjacent to each other (e.g. C_3_-A_1_, C_3_-B_1_, A_3_-D_1_, etc.). *Within* similarity was significantly higher than *across* similarity for duplet pairings. (*Median difference* = 0.03, *V*(53) = 1147261242, *p* < 0.0001, *Bonferroni corrected, CI* = [0.03, 0.03], Wilcoxon rank sum test). (**Fig. 4**)

**Figure 4.**
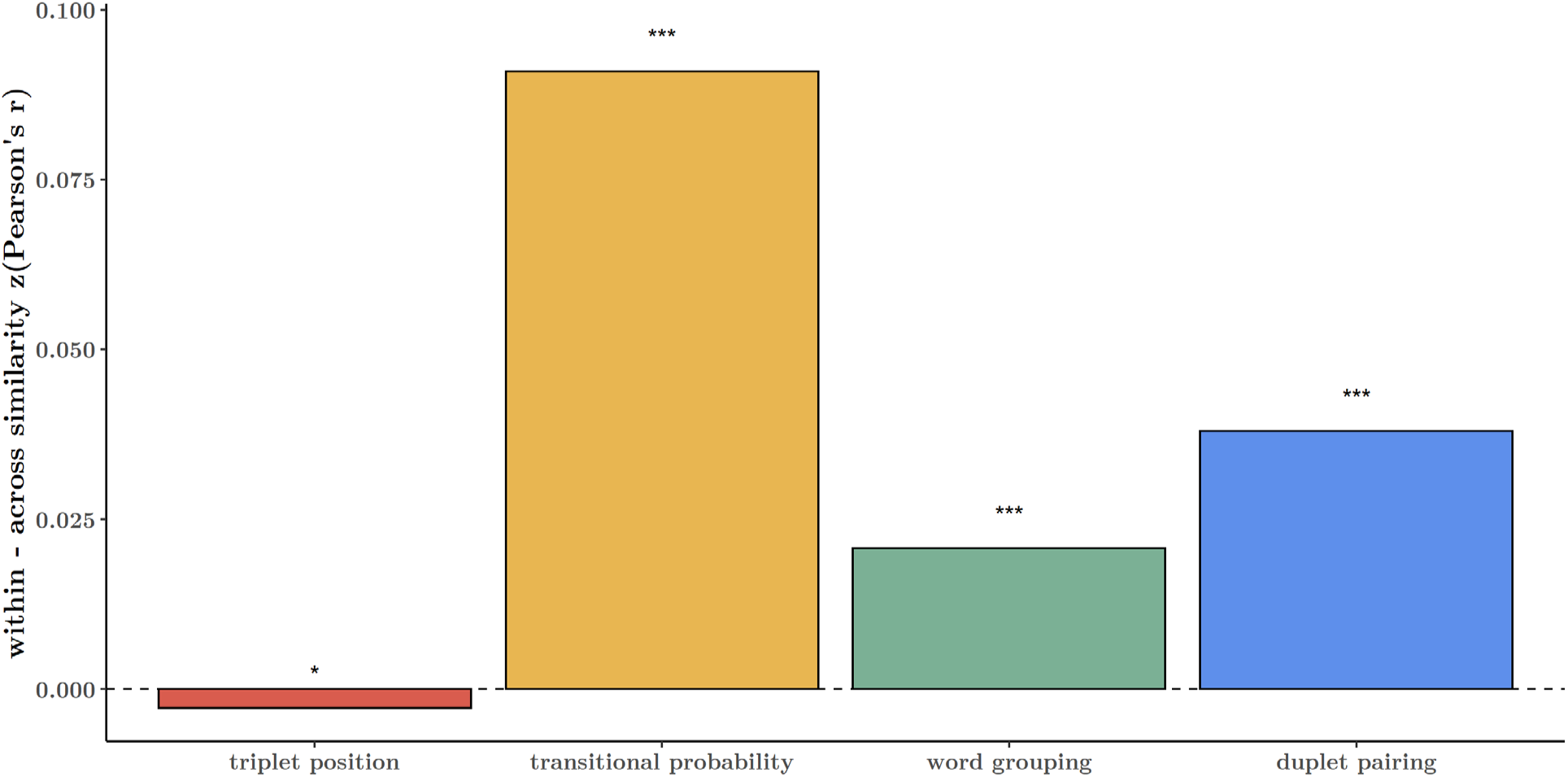
RTs reveal representational similarity for triplet position, transitional probability, word grouping, and duplet pairings. Representation similarity analysis showed that groups of RTs classified as belonging to the within category were more similar than groups of RTs classified as the across category for features transitional probability, word grouping and duplet pairings (within a given pseudoword). Across similarity was greater than within similarity for feature triplet position. (Wilcoxon’s rank sum test for paired groups, on bootstrapped z-transformed Pearson’s correlations between syllables for each participant.) *p<0.05, ***p<0.0001

#### Sensitivity to Transitional Probability Weakly Predicts Word Recognition Performance

Finally, we wished to test whether there exists a correlation between word recognition performance and the similarity measures derived from the RSA. We repeated the RSA at the subject level with only those participants from Experiment 1 who completed both online and offline tasks (N = 32). For each participant, we obtained a measure of similarity (*within-across* median correlations) for each of the four features. These similarity measures were then correlated with each participants’ word recognition performance. We observed virtually no relationship between word recognition accuracy and *within-across* median similarity for transitional probability (*ρ* = 0.02, *t*(30) = 0.12, *p* = 1,*Bonferroni corrected*), triplet position (*ρ* = 0.03, *t*(30) = 0.19, *p* = 1,*Bonferroni corrected*), word grouping (*ρ* = −0.2, *t*(30) = −1.14, *p* = 1,*Bonferroni corrected*), or duplet pairing (*ρ* = 0.18, *t*(30) = 0.1, *p* = 1,*Bonferroni corrected*, Pearson’s product-moment correlation, one-sided). (**Fig. 5**)

**Figure 5.**
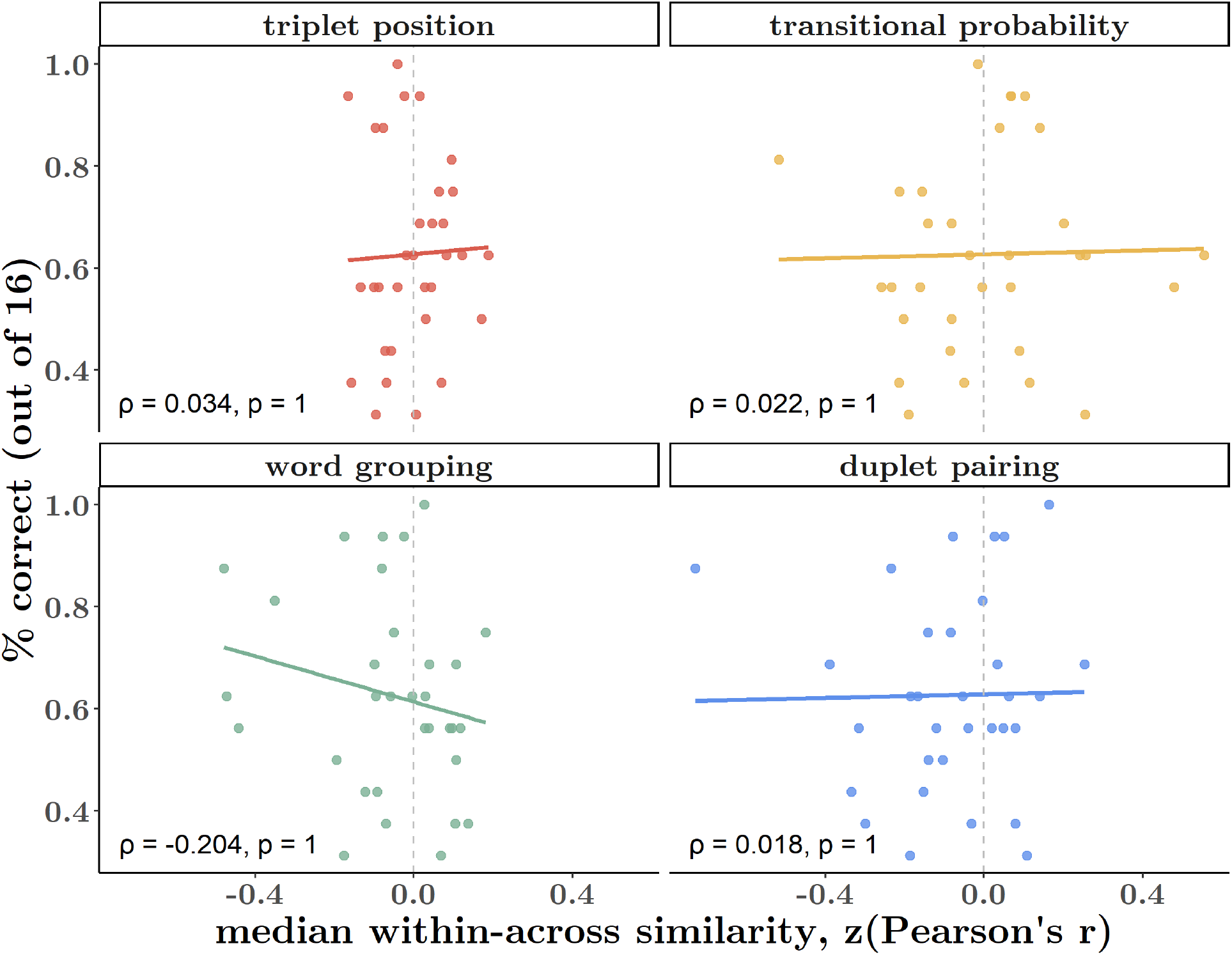
RT similarity measures do not correlate with word recognition performance. Word recognition performance (% correct trials) does not correlate strongly with similarity (measured as within-across median RT correlations) for any of the four features tested: triplet position, transitional probability, word grouping, or duplet pairing. (One-sided t-test on Pearson’s product-moment coefficient.)

#### Discussion

Using RSA, we aimed to identify the specific features of the speech stream for which target detection tasks evaluate learning. Our RSA revealed that RTs carry information about multiple features of the syllables: transitional probability, triplet position, word grouping, and duplet pairing of syllables. This result indicates that participants are specifically sensitive to pairwise relationships between syllables, as well as the triplet sequence.

*Within* similarity values were higher than *across* similarity values for transitional probability, word grouping, and duplet pairing. For transitional probability, this may be due to the fact that the higher predictability of *within-*syllables led to more consistent RTs, as compared with RTs to the less predictable *across* or word-initial syllables. For word grouping, this result may suggest that the pattern of RTs for syllables comprising a pseudoword is somewhat unique to that particular sequence of syllables (i.e. the degree of facilitation of that specific word-initial syllable on the detection of the following word-medial syllable, and the latter’s facilitation of the word-final syllable). For duplet pairings, we also found that RTs are more similar for syllable pairs within pseudowords than for pairs that cross word boundaries. This is to be expected, as there exists no strong relationship between word-final syllables of one pseudoword and word-initial syllables of another.

However, similarity *across* triplet positions was higher than similarity *within triplet positions.* At first, this may seem surprising, as *within* pairs share the same ordinal position within pseudowords. However, *across* pairs include many pairs that are within-word duplets, as well as phantom word pairs (e.g. C_2_-D_3_) that share the same transitional probability, whose similarity to each other may together override the similarity between syllables of the same position within a pseudoword. This finding underscores strong influence of pairwise relationships in driving RTs.

Finally, by correlating subject-level similarity values for each feature with participants’ word recognition scores, we aimed to pinpoint whether sensitivity to these features is shared between these two tasks. However, we found there was virtually no correlation between word recognition and similarity values for any of the features tested. While we cannot infer that these features were not instrumental in helping participants complete the word recognition task, we might conclude that the word recognition task does not (in our instantiation) have sufficient granularity or power to predict performance on online SL tasks.

## General Discussion

Statistical learning is an impressive cognitive tool: it is fast, flexible, and robust. It occurs automatically [13] and independent of input modality or stimulus type [31]. Yet, this seemingly simple mechanism can be challenging to evaluate accurately, as it encompasses several learning and memory components, including stimulus encoding, retention, and abstraction. [32],[33] As a field, we require sensitive and precise tools to discern (1) what information is being extracted by participants during exposure, (2) what of this information they are able to exploit in behavioral tasks, and (3) what processes or knowledge are being measured by our tasks. Given the richness of the SL process, an adequate evaluation of SL ability likely requires the use of multiple measures within a given study, as well as a thorough understanding of how these measurements relate to one another, i.e. what they jointly or distinctly measure.

We showed that online target detection tasks are highly sensitive, multi-faceted tasks, with the potential to provide insight into the contribution of distinct structural features to behavioral responses. We were also able to replicate above-chance word recognition performance, using a canonical 2AFC pseudoword vs. part-word task. Intriguingly, although we observed a weak correlation between performance in these two tasks, RT similarity measures derived using RSA were unable to predict word recognition performance for any of the four features tested.

Previous proposals noting the lack of correlation between canonical online and offline tasks have pointed to the different demands the tasks make on memory, [20],[24], and the psychometric strength and weakness of the measures, respectively, [12] to explain the discrepancy. Indeed, in our study, above-chance performance was present in only 71% of participants (of 33) in the 2AFC task, while the graded RT effect was present in 89% of participants across both experiments (of 53). (**Fig. S7**)

As discussed above, the 2AFC task is a double-edged sword: too many trials risks reinforcing memory for non-words or part-words and obscuring the subtle contribution of implicit statistical learning, while too few trials provide a noisy and underpowered measure of learning. Consistent with previous studies [21],[34], we tested each pseudoword 4 times, in an attempt to minimize the former issue. However, the optimal tradeoff between these issues likely resides in a test where a larger number of test items (e.g. pseudowords) are tested with few repetitions (e.g. 4-8), yielding a minimum of 32 or 36 trials.

In our study, the interpretability of our results are constrained by covariance among the features of interest, which disallows us from making firm assertions as to which features were more or less instrumental than others in completing the task. Future studies can make maximal use of the sensitivity of online tasks and RSA by incorporating designs that de-confound as many distinct properties as is reasonable or desired, to tease apart the relative contributions of e.g. transitional probability and word grouping or co-occurrence frequency [8].

### Task sensitivity and its theoretical implications

Saffran et al.’s seminal study concluded with the suggestion that infants exposed to the continuous syllable stream “succeeded in learning and remembering particular groupings of three-syllable strings.” [5] Since this study, which catalyzed interest in SL reach in the domain of language, there has been debate over how participants accomplish the task. While there is evidence for numerous forms of sequence learning in general (such as rule-based learning that allows the learning of non-adjacent relationships [1],[35]), there are two accounts that seek to explain Saffran-type SL tasks.

On what may be called the “chunking” or memory-based view, memory of the word chunk is what allows successful performance in the 2AFC pseudoword vs. part-word discrimination task. [33],[36],[37] In the PARSER model of chunking by Perruchet & Vinter (1998) [36], the perceptual system is thought to generate chunks of arbitrary length (1, 2 or 3 syllables) from the input, which are then weighted (selected for or against as meaningful units) by repeated exposure to the same template. They propose that the chunk is subsequently operated upon as a single unit. Given the emphasis on the formation and processing of the chunk as a whole, this characterization does not appear to support the observation that predictability for elements within the chunk may vary.

In contrast, the simpler process of tracking local transitions between items predicts the common observation that in, e.g. a syllable triplet, initial syllables predict medial syllables, which in turn predict final syllables. [38] We note however that these explanations are not mutually exclusive, as all different forms of sequence processing are available to the individual. In artificial languages with non-adjacent dependencies [39], or more enriched syntax [40], more complex, hierarchical tracking of inputs would be required. [41]

Importantly, success in both the online and offline tasks does not require nor entail that an individual was able to generate a unified representation of the syllable triplets that formed the pseudowords. Sensitivity to transitional probabilities (mere segmentation) may be sufficient to achieve both the graded RT effect and above-chance word recognition. As discussed above, learning the pairwise transitions between syllables will facilitate responses to word-medial and word-final syllables.

Likewise, in the word recognition task, the sum transitional probabilities for isolated pseudowords is 2 (TP = 1 between syllable 1 and 2, and also 1 between syllable 2 and 3), but only 1.33 for part-words (TP=0.33 between syllable 1 and 2, and 1 between syllable 2 and 3). While we are agnostic as to how precisely participants complete the tasks, those used in this study can be minimally explained by learning of a single feature.

The notion that tracking transitional probabilities alone would allow above-chance performance on the pseudoword vs. part-word recognition task, but not be sufficient for “chaining” or “chunking” more than two items is also supported by Endress & Mehler’s findings. [10] In their study, 5 minutes of exposure was sufficient for participants to discern words from part-words, but even after 40 minutes of exposure, they still could not discern words from phantom words (tri-syllabic sequences where syllables are drawn from different words, but ordered so as to maintain each syllable’s original position in the word). Only with the introduction of linguistic cues (final syllable lengthening and pauses between words), did participants reject phantom words as often as they rejected part-words without additional cues.

Another study found that correct judgments of whether a novel triplet belonged to an exposure stream of repeating visual shapes were improved by how closely the summed transitional probabilities within the new triplet matched that of the original triplet structure. [42] Finally, a recent study specifically compared participants’ ability to learn sequences of two versus three items, finding that learning triplet sequences required explicit instruction, while duplet sequences could be acquired implicitly. [43]

Our study highlights the importance of implementing complementary measures of SL within the same experiment with adult participants. However, one implication of our results is that online tasks may in some cases and for some experimental questions be preferable to explicit, discrete 2AFC tests. This is particularly relevant for SL studies on infant populations. Gaze detection tasks, common in infant research, more closely approximate the 2AFC task. Indeed, for infants and young children it is more challenging to find an analogue to the online target detection task.

However, studies in infants have found rich, complementary evidence for learning by utilizing frequency tagging techniques during exposure [44], as well as measuring ERP components [44],[45] or pupilometry [45] during the test phase. Frequency tagging [7], ERPs [21] and pupilometry [46] are all methods that have also revealed neural correlates of learning in adults.

The rapid and implicit structure learning exhibited by humans and other non-human animals is impressive in its flexibility and domain generality, easily spanning vision, audition, and motor action. [47] What we call SL is therefore likely to be a complex collection of several learning systems for syntactic tracking, memory encoding and retrieval, and hierarchical processing. [31],[48]

## Conclusion

Online measures of statistical learning, such as the target detection task, can reveal subtle, dynamic properties of implicit learning. Meanwhile, offline tasks that require explicit discrimination show that participants can use implicitly acquired knowledge to make explicit decisions, but are limited in their inferential power. We found evidence of learning using both tasks, but found a weak correlation between these measures. Using representational similarity analysis, we found that RTs to target syllables in a continuous syllable stream captured information about the syllable’s transitional probability, triplet position, word grouping and duplet pairing. Furthermore, RT similarity measures for these features were unable to predict word recognition performance. We conclude that online tasks appear to capture more subtle information than typical explicit recognition tasks, even in less-than-ideal experimental designs. We highlight the importance of using multiple methods to assess SL, developing thorough theoretical models, and subtle tasks to investigate this sophisticated and multi-component ability.

## Methods

### Stimuli

Speech stimuli consisted of 12 consonant-vowel (CV) pairs. We selected 5 unique vowels that are maximally separated in their manner and place of articulation. We ensured that none of these vowels typically occurred in unstressed syllables in spoken German. We then selected 12 unique consonants, in order to render each syllable phonetically distinct from the others. We used the CELEX database [49] to calculate the frequency of occurrence of each of our syllables in spoken German, as well as the frequency of co-occurrence between each pair of syllables. We eliminated high-frequency CV pairings from our list of possible syllables and formed the final words by combining three syllables (each with distinct vowels) for which no transitions were frequent in spoken German. The syllabes were: be, di, ga, ki, la, mi, nu, po, ro, se, tu, za.

A male native speaker of German was recorded pronouncing each syllable in our set separately and with a flat intonation. Each syllable was repeated several times to ensure we obtained a quality token. The token which most closely followed the IPA pronunciation was selected as the final syllable. The syllables were then high-pass filtered at 50 Hz and silences before and after syllable were removed using a custom script in Matlab 2017b. The 12 syllables were normalized for pitch and intensity using Praat [50] to ensure relative homogeneity between tokens. Finally, syllables were temporally compressed to 240 ms in duration and a 10 ms silence was added at the end of each syllable, for a total duration of 250 ms.

Syllables were combined into 4 tri-syllabic pseudowords such that each word featured no repeating consonants or vowels and similarity between any possible succeeding pairs of syllables was minimized. We also ensured that no pairs were phonotactically illegal or shared a resemblance with existing words in German, using CELEX. Pseudowords for our study were: nugadi, rokise, mipola, zabetu. Part-words, used in the word recognition task in Experiment 1, were of the form C’AB (word-final syllable from one word followed by word-initial and word-medial syllables from another): dizabe, semipo, lanuga, turoki.

#### Experiment 1

Continuous speech sequences (24) were created in Matlab by concatenating syllables comprising the four pseudowords such that no words repeated consecutively. Each stream was comprised of 216 syllables (72 words) and was 54 seconds long. As per the design in [5], standard in SL studies, the only cue to segmenting the sequence lay in the transitional probabilities between syllables. The transitional probability of word-medial and word-final syllables (relative to the preceding syllable) was 1, while the transitional probability of word-initial syllables was 0.33. The first syllable in each stream could be a word-initial, word-medial, or word-final syllable. If stream began with the word-medial (word-final) syllable of a word, the word-initial (word-initial and word-medial) syllable of that word would be the last (two) syllable(s). Speech streams were ramped up and down in amplitude using a linear slope over a period of 1.5 seconds (6 syllables) so that onset and offset syllables were not clearly distinguishable and could not serve as cues to word segmentation.

#### Experiment 2

For this experiment we synthetized 12 “structured” streams and 12 “random” streams in Matlab. For structured streams, the procedure was identical to that mentioned above. For random streams, the 12 syllables were pseudo-randomly permuted out to the same length as the structured stream (216 syllables), with the sole constraint that a syllable could not be repeated consecutively. Thus, transitional probabilities between adjacent syllables were roughly 0.083. Speech streams were ramped up and down in amplitude over a period of 1.5 seconds so that onset and offset syllables were not clearly distinguishable and could not serve as cues to word boundaries.

### Procedure

The experiment was designed using Presentation^®^ (Version 20.1 Build 12.04.17) and delivered on two versions of the software (Version 20.0 Build 07.26.17 and Version 21.1 Build 09.05.19, Neurobehavioral Systems, Inc., Berkeley, CA, www.neurobs.com).

#### Experiment 1

41 individuals participated in the study (27 female, mean age, 27.44 ± 5.78 sd). Participants provided written informed consent prior to the study. The study received approval from the local ethical committees (Ethics Council of the Max Planck Society) and adhered to the ethical standards of the Declaration of Helsinki. All participants reported having normal hearing and were paid for their participation. Two participants were removed from the data pool due to technical failure. Of the 39 remaining datasets, 33 were used in analyzing the target detection task (one participant failed to follow instructions, and technical issues caused partial data loss for the other five). Since the design of our experiment was modular, technical failure in one task did not necessarily affect data in another. Of the 39 datasets, we were able to use 38 for analyzing the word recognition task (data from one participant in this task was overwritten). For the correlation analysis comparing target detection task performance with word recognition performance, we included only participants for whom we had data for both tasks (32).

A previous study by Batterink and colleagues [21] using similar online and offline tasks as us had observed a significant correlation coefficient of 0.51 with 24 participants. A power analysis revealed this analysis to have a power of 0.74, suggesting that this effect size is rather large based on Cohen’s effect sizes for *ρ* values of 0.1, 0.3, and 0.5, respectively representing small, medium, and large effects. We calculated that in order to obtain a test with at least 80%, we would need 27 participants, and for 90% 36 participants. Our sample of 33 then was theoretically sufficient to observe a correlation effect as large as Batterink et al. reported.

Participants were seated in a dimly-lit, sound-attenuated booth, approximately 52 cm from the monitor and listened to the stimuli via headphones connected to a headphone amplifier (Beyerdynamics-DT-770 80 Ohm; Lakepeople G103P1262). Stimulus intensity level was approximately 57 dB (LAF: min 44 dB, max 76 dB), as measured by a NTi Audio device connected to an artificial ear on which the experiment headphones were mounted. The experiment was conducted on a 64-bit Windows machine (Fujitsu Celsius M740B) running Windows 10.

The experiment consisted of an exposure phase, during which participants performed the target detection task, followed by the word recognition task. Our experiment also included an additional task, designed to measure perceived speed of the speech stream before versus after the exposure phase. Results from this task will not be discussed here.

During the exposure phase, participants listened to a total of approximately 24 minutes of continuous speech. Participants were told they would hear brief sequences of sounds from an alien language. Audio was presented binaurally. Before the start of each stream, one of the 12 syllables was displayed orthographically on the screen and played aurally twice. Participants were instructed to press the spacebar as fast as they could during the subsequent stream whenever they heard this target syllable. Each of the 12 syllables served as a target syllable twice. The presentation order of syllables was pseudo-randomly shuffled for each participant with the constraint that a syllable from each triplet position in the pseudoword (word-initial, word-medial, or word-final) was tested before any were repeated. The 24 streams were organized into 8 blocks, where each block consisted of 3 streams with one target syllable from each triplet position tested. Within each stream, target syllables appeared 17-18 times. Participants could take self-paced breaks between blocks.

In the word recognition task, participants completed 16 trials of a two-alternative forced-choice task. In each trial, a pseudoword and a part-word were presented (counterbalanced across trials), and participants were prompted to determine which of the pair was a word in the alien language they had just heard in the previous section. The inter-stimulus-interval between words was 400 ms, while inter-trial-interval was 1.2 seconds. Each pseudoword was paired with each part-word once (4 x 4 trials).

#### Experiment 2

21 individuals participated in the study (15 female, mean age 28.08 ± 6.82 sd). Participants provided written informed consent prior to the study. The study received approval from the local ethical committees (Ethics Council of the Max Planck Society) and adhered to the ethical standards of the Declaration of Helsinki. All participants reported having normal hearing and were paid for their participation. Inclusion criteria included the requirement to not have taken part in Experiment 1. One participant was excluded due to technical failure. Technical failure caused data loss in the random condition for one other participant, leaving data from 20 participants (19 in the random condition, 20 in the structured condition).

Participants were seated approximately 52 cm from the monitor and listened to the stimuli via headphones connected to the PC server. Stimulus intensity level was again measured by a NTi Audio device connected to an artificial ear. Volume levels were in the range reported for Experiment 1. The experiment was conducted on a 64-bit Windows machine running Windows 7.

Participants completed two exposure phases, one with a continuous stream of random syllables and one with a continuous structured stream. During both phases, participants completed the target detection task, following the procedure described under Experiment 1. Each phase consisted of a total of approximately 12 minutes of continuous speech, divided into ~1 minute long streams. Participants could take self-paced breaks between streams. The instructions and task procedure for each phase was identical to that in Experiment 1, with the exception that participants only performed the task once for each syllable instead of twice. Each stream featured 18-19 occurrences of the target syllable. Random and structured exposure orders were counterbalanced across participants. Our experiment also included an additional non-SL task, which was completed after each exposure phase. Results from this task will not be discussed here.

### Analysis

All analyses were performed in RStudio (version 1.2.1335; RStudio Team 2018) using the R statistical programming language. [51] Primary analyses and modelling was performed using the packages *tidyverse* [52]*, lme4* [53], and *emmeans.* [54] Raw data was transformed into csv files for processing in R using Matlab R2017b (version 9.3.0.713579). Data and code is publicly available on Github: https://github.com/avakiai/statistical-learning.

#### Experiment 1

Online Target Detection Task: We considered only those responses that occurred within a boundary of ± 3 times the median absolute deviation over all RT values. This procedure ensures that RT cutoffs would be based on the distribution of the raw data and not arbitrary limits [55]. At the same time, the use of the median as the centrality metric is arguably more appropriate, given that the mean can be a biased estimator of RT data, which typically follows a gamma, lognormal, or ex-Gaussian distribution. This procedure eliminated only 0.034% of the data and resulted in RT that ranged from 0 to 943 ms (versus the original 0 to 1298 ms). This procedure did not significantly change the overall mean accuracy (*M*_*before*_ = 0.71, *SD*_*before*_ = 0.45, *t*(63.9) = −0.39, *p* = 0.7, two-sided t-test against post-outlier removal accuracy).

To replicate findings that showed graded reaction times in response to syllables in different triplet positions, we ran a generalized linear model with RT (in seconds) as outcome variable, fitted with a gamma distribution and log link function. Our fuller model included both triplet position and block as fixed effects factors, and subject as a random intercept-random effects factor. This model was compared with a lesser model in which only triplet position was used as a fixed effect. The lesser model provided a better fit of the data, with a lower AIC (−6389.2) value and significantly lower deviance (−6399.2, *X*^2^(21, *N* = 33) = 49.066, *p* < 0.001). (See **Table S1** for regression results.) We also compared both the fuller and the lesser models with random slopes for levels of triplet position in the random effects term, but the lesser model with only varying random intercepts in the random effects term still proved a better fit for observed data (see **Table S1;** lesser vs. fuller random slopes models deviance was −6624.3 and −6678.3, respectively; *X*^2^(21, *N* = 33) = 54.025, *p* < 0.0001). Thus, we conducted further analysis on results of the lesser model.

Word Recognition Task: We computed each participant’s word recognition accuracy by dividing the number of correct discriminations by total trials (16) for each participant. Overall mean recognition accuracy was the overall number of correct divided by total trials. Recognition accuracy per word was calculated in the same manner, but for each word individually. Chance performance was set at 50% correct trials (8 out of 16 for overall accuracy, 2 out of 4 for word-wise accuracy).

Correlation between online and offline measures of SL: Three methods were used to evaluate the correlation between performance in the two tasks. For the analysis following Siegelman et al. [26], the online measure of SL was computed for each participant (see equation 1) and correlated with participants’ word recognition accuracy scores using Pearson’s correlation (one-tailed test). For the analysis following Batterink et al. [21], an RT score was created for each participant computing RT difference between word-initial and word-final position syllables. These scores were correlated with word recognition scores using Pearson’s correlation. Finally, we used a third method in which we z-normalized RT values for each participant, computed median RTs to each triplet position, and computed the difference between the scaled median RTs for each position pairing for each participant (1-2, 2-3 and 1-3). We then correlated each of the three RT scores with word recognition accuracy separately using Pearson’s correlation.

#### Experiment 2

Online Target Detection Task: We used the same criterion to eliminate outliers as in Experiment 1 (± 3 times the median absolute deviation). This procedure eliminated only 1.93% of the data and resulted in RT that ranged from 119 to 941 ms (originally, 0 to 1997 ms). This procedure did not affect the overall detection accuracy (*M*_*before*_ = 0.83, *SD*_*before*_ = 0.38, *t*(37.43) = −0.42, *p* = 0.68, two-sided test against post-outlier removal accuracy)

To compare the reaction times between structured and random conditions, we dummy-coded the random streams with the same triplet positions as the structured streams. Thus, if the syllables in the structured stream 1 followed the order: 3,1,2,3,1,2,3…, we applied the same position coding to random stream 1, even though these position codes correspond to no meaningful property in the random stream. This procedure however, allowed us to compare RTs for the same variable (triplet position) between the two conditions.

We performed a modelling procedure similar to that from Experiment 1. We included subject as a nested effect within condition order (whether participants completed the structured condition before the random condition, or vice versa), as condition order was our between-subjects variable. We further specified the random effects term by allowing random intercepts and uncorrelated random effects for each level of condition. This structure allows the graded RT curve for each participant to vary between conditions, as well as their baseline RT (intercept). (See **Table S2** for regression results.)

#### Representational Similarity Analysis

For the group-level analysis, we first combined the preprocessed data (after outlier removal) from Experiment 1 (N = 33) with the data from the structured condition in Experiment 2 (N = 20) for a combined data set with greater power (N = 53). For each participant, we computed a similarity matrix (Pearson correlation) on RTs between each pair of syllables, thus generating a 12-x-12 matrix of correlation values for each participant. We then applied a Fisher’s z-transformation to each matrix to normalize correlation values across participants. We tested for the coding of each of the four features by running four Wilcoxon ranked sum tests. For each test, we identified the cells in the 12-x-12 matrix whose values would comprise *within* and *across* groups, respectively. We performed a random sampling of correlation values from the cells in the matrices from all participants to generate *within* and *across* arrays. (See Results and **Fig. S6**.) The sampling procedure was repeated 200 times with replacement, with the final N for each test being equal to 4/5 times the length of the shorter of the two arrays being compared. (Since arrays for paired ranked sum analyses must have the same length and not all tests entailed the same number of comparisons, we effectively subsampled from both arrays so they would have a common length.) We then computed a paired, two-sided Wilcoxon’s rank sum test on the resulting two arrays to determine whether similarity (as a proxy for feature coding) is higher within or across groups.

For the subject-level analysis, we took the z-transformed correlation matrices from participants in Experiment 1 (N = 32) and for each participant individually, subset values from the matrix to generate within and across groups for each feature, using the same rubric as in the previous analysis. (However, here we did not perform a bootstrapping procedure.) Next, for each feature, we computed mean of the correlation values in the within and across groups, and subtracted the across mean from the within mean, to obtain four measures of similarity for each participant – one for each feature. Finally, we correlated each participant’s word recognition score with their within-across similarity for each feature individually, to determine if sensitivity in the RT task to any of these feature alone would predict word recognition performance.

## Acknowledgements

We thank Dr. Yue Sun for help with stimulus creation. We also thank Drs. R. Muralikrishnan & Cornelius Abel, as well as Hanna Kadel & Dominik Thiele for technical and administrative support. Many thanks go to Valeria Peviani, Martina Vilas, Alex Lepauvre, & Valerie Pu for helpful comments and camaraderie. This research was supported by the Max Planck Society and the German-Israeli Foundation for Scientific Research and Development to L.M.

## Contributions

A.K.: Conceptualization, Methodology, Investigation, Software, Formal Analysis, Visualization, Writing – original draft and review & editing. L.M.: Conceptualization, Methodology, Supervision, Funding Acquisition, Writing – review & editing.

