## Supplementary Materials for "What canonical online and offline measures of statistical learning can and cannot tell us"

### Supplementary Information

#### *Experiment 1*

<u>Detection accuracy</u>

We examined detection accuracy overall (See Results), as well as a function of ordinal position, pseudoword, and target syllable, to ensure that difference in the stimuli did not drive our main effects.

We found that detection accuracy was modulated by ordinal position ($F(2) = 24, p < 0.0001$), with each successive position having a higher mean accuracy ($M_{position\ 1} = 0.59, sd = 0.49$; $M_{position\ 2} = 0.70, sd = 0.46$; $M_{position\ 3} = 0.82, sd = 0.39$). Pairwise contrasts revealed mean differences between RTs to each position to be significant ($1-2{:}\ t(96) = -3.41, p = 0.003, Cohen's\ d = -0.84; 1-3{:}\ t(96) = -6.93, p < 0.0001, Cohen's\ d = -1.7; 2-3{:}\ t(96) = -3.52, p = 0.002, Cohen's\ d = -0.87$).

Accuracy also varied between the four pseudowords ($F(3) = 11.53, p < 0.0001$). Specifically, between words *nugadi* and *zabetu* ($t(128) = 5.52, p < 0.0001, Cohen's\ d = 1.356$), *rokise* and *zabetu* ($t(128) = 4.15, p = 0.0003, Cohen's\ d = 1.02$), and *mipola* and *zabetu* ($t(128) = 4.25, p = 0.0002, Cohen's\ d = 1.04$). (**Table S3a**) This suggests that the accuracy was overall equal between all words except *zabetu*, for which detection was more difficult.

Finally, we found that syllable identity also affected detection accuracy ($F(11) = 20.38, p < 0.0001$). Certain CV syllable pairs may have been easier to detect than others, due to minor variations in stimuli acoustics. (**Table S3b, Fig. S1a**) We interpret these results as meaning that (1) the predictability of syllables in positions 2 and 3 made their detection more likely than syllables in position 1, and (2) that certain syllables were more easily detected, which may have led to the difference in detectability at the pseudoword level (which combines the detectability of the three syllables which comprise it). We addressed these two possible confounds in two analyses.

<u>Detection accuracy and RT</u>

First, to ensure that the difference in detection accuracy between ordinal positions did not affect the results, we ran the same model we used for our primary analyses (see results for Experiment

1) with a subsample of RT data. Since accuracy varied between positions, the number of observations for each levels of factor ordinal position varied also. We subsampled our data set so that the observations for all ordinal positions were equal to the smallest group ($N_{position\ 1} = 2{,}654$). We observed the same main effect of ordinal position ($X^2(2, N = 33) = 458.8, p < 0.0001, \text{Type II Wald Chisquare test}$). All pairwise contrasts between ordinal positions reached significance ($p < 0.0001$). Given this result, and the fact that generalized mixed models are robust to unbalanced data sets, we concluded that the unequal number of observations between position levels did not skew our results.

*Regressing out variability in individual syllables in RT effect*

Secondly, we sought to validate the results discussed in the main text by regressing out the effect of individual syllable as a function of ordinal position. We ran a generalized mixed model with ordinal position and target syllable as fixed effects factors and subject as random effect factor, with the RT (in ms) as outcome variable. We then subtracted the resulting residual values for each data point from the raw RT, and re-ran the lesser model as specified above using the adjusted RT values. We still observed the main effect of ordinal position ($X^2(2, N = 33) = 538.53, p < 0.0001, \text{Type II}$) and significant pairwise contrasts between all three levels of ordinal position ($p < 0.0001$). Therefore, we concluded that slight variations in the acoustics of our stimuli did not significantly affect our results.

*Contra a trivial graded RT effect*

To address the critique put forward by Himberger et al., that a confound of stream and triplet position, combined with a general facilitation of RT over the course of the test, could trivially generate an RT effect, we performed a few additional analyses. For this critique to be valid, the mean difference in RT to targets in positions 1 & 2 and 2 & 3 must be equally large, and RTs must decrease over the course of exposure (both within trials and across blocks). To address this confound, we first performed a linear regression on mean RT differences between each position pairing (1-2, 2-3, and 1-3) with both block and pairs as predictors.

We found no interaction between block and pairs, suggesting that the magnitude of the differences between positions, as well as the relationships between them, did not change over the course of the blocks ($X^2(14, N = 33) = 5.93, p = 0.97$). We found only a main effect of pairs ($X^2(2, N = 33) = 28.42, p < 0.0001$, Type II). (**Fig. S2a-b**) Tukey-corrected contrasts between pairs showed the mean differences between position 1-2 ($M_{position\ 1-2} = 59\ ms, SD = 135\ ms$) were significantly larger than those between positions 2-3 ($M_{position\ 2-3} = 29\ ms, SD = 120\ ms$; $t(613) = 2.71, p = 0.019, Cohen's\ d = 0..24$), supporting the notion that the RT effect is not linear between the positions. Since the decrease in RT observed between positions 1 & 2 is indeed larger than that between positions 2 & 3, the effect cannot be due to a simple confound of monotonically decreasing RTs over the course of the session.

Additionally, there was no effect of block on overall RT ($X^2(7, N = 33) = 8.99, p = 0.25$, Type II) nor any difference between RTs when looking specifically at blocks 1 and 8 ($t(49.93) = -0.60, p = 0.55$)). Furthermore, we graphically demonstrate that the distribution of target syllables in all triplet positions is uniform across stream positions (1-216). (Plots for Experiment 1 stream positions shown. **Fig. S2e-f**)

#### *Experiment 2*

<u>*Detection accuracy*</u>

In Experiment 2, we tested for an effect of condition, ordinal position, pseudoword, and syllable identity on detection accuracy, to ensure the reliability of our results. Note that pseudoword and ordinal position are only meaningful factors in the structured condition. As we report in the main text, detection accuracy in the structured condition was numerically higher than in the random condition, but this difference did not reach significance at the 5% alpha level.

We found no difference in accuracy as a function of ordinal position in the structured condition ($F(3) = 0.79, p = 0.46$). We also found no difference in accuracy as a function of pseudoword in the structured condition ($F(3) = 0.93, p = 0.43$). Next, we checked for an effect of syllable identity in both conditions. We found no interaction between condition and target identity, but main effects of both on detection accuracy ($target\colon F(11) = 3.35, p = 0.0001; condition\colon F(1) = 9{,}27, p = 0.002$). We computed pairwise contrasts between target syllables, collapsed over both conditions. Only a few contrasts reached significance: *ga-di, di-be, ki-be, se-be, po-be* (all p < 0.05). This would suggest the effect is mainly driven by lower detectability for a few, specific syllables, e.g. *be*. (**Fig. S1b, Table S4**)

<u>*Regressing out variability in individual syllables in RT effect*</u>

To be confident in our results, we performed two follow-up analyses to ensure that the variability observed in detection accuracy for certain syllables did not bias our results. We validated our results by regressing out the effect of individual syllables as a function of ordinal position. We ran a generalized mixed model with ordinal position and target syllable as fixed effects factors and subject as random effect factor, with RT (in seconds) as outcome variable. We then subtracted the resulting residual values for each data point from the raw RT, and re-ran the lesser model (see results for Experiment 2) using the adjusted RT values. We still observed the main effect of ordinal position ($X^2(2, N = 20) = 354.08, p < 0.0001, \text{Type II}$) and significant pairwise contrasts between all three levels of ordinal position ($p < 0.0001$). Therefore, we concluded that, once again, minute differences between stimuli did not affect our main results.

<u>*Carry-over effects of structure, random condition on RT*</u>

We additionally wished to confirm that there were no carry-over effects of condition, where the order in which participants saw the two conditions of the target detection task might influence the RT effect we were aiming to replicate. We ran a generalized linear mixed model with RT (in seconds) as outcome variable and condition, ordinal position, and condition order as predictors. We set the random effects term as a random intercept term with subject nested within condition order. We observed no three-way interaction between our three predictors ($\chi^2(2,\ N = 20) = 0.33,\ p = 0.85$), suggesting that condition order did not significantly bias our results. However, we do observe a difference in the plotted median RT values when comparing structured and random condition in each of the two condition orders. (**Fig. S5**) It is plausible that a minor carry-over effect did occur, but our analysis was too underpowered to detect it.

### Supplementary Figures

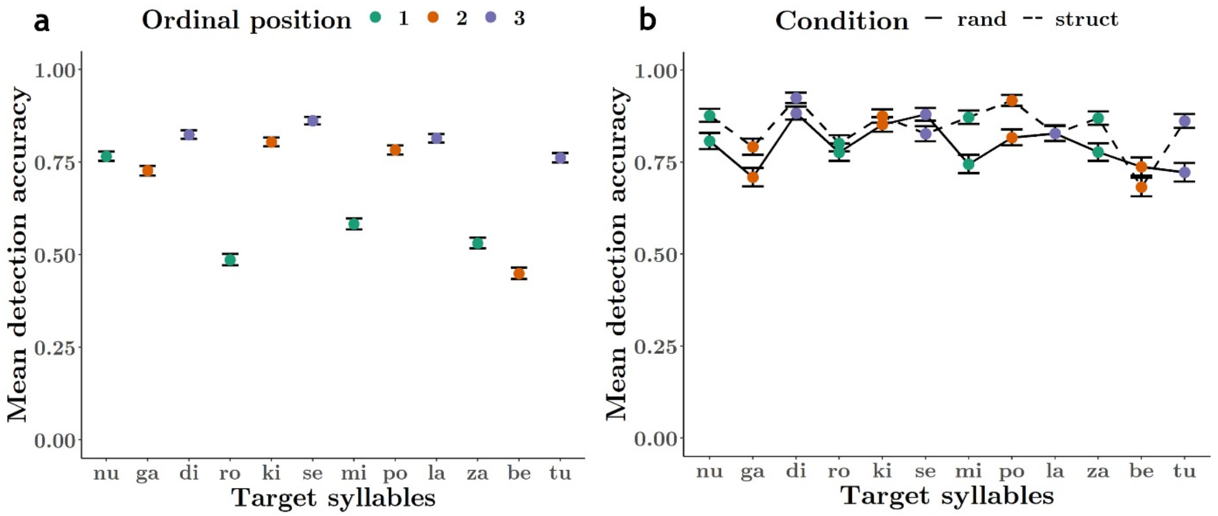

***Figure S1. Detection accuracy in Experiments 1 & 2.*** *A. Overall detection accuracy (hit rate) for each target syllable in Experiment 1. Colors indicate ordinal position of each syllable. Accuracy was generally lower for 1st position syllables vs. 2nd and 3rd position syllables. B. Overall detection accuracy (hit rate) for each target syllable, in each condition, in Experiment 2. Error bars represent SEM. Accuracy was higher for target syllables in the structured condition versus the random condition.*

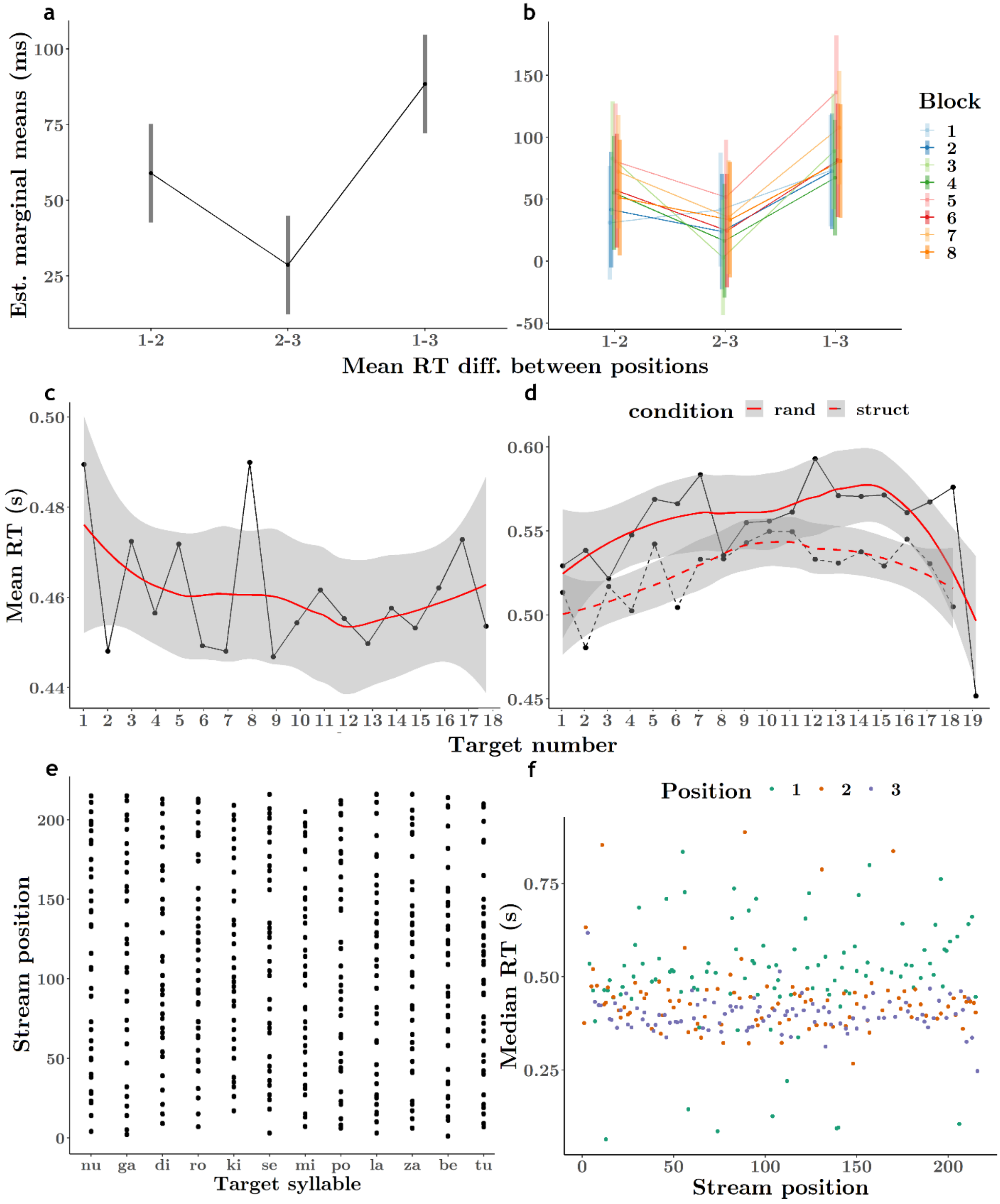

***Figure S2. Exp. 1: Controls to disconfirm effect of spurious RT facilitation.*** *A. The mean difference in RT between 1st and 2nd position targets was higher than the mean difference between 2nd and 3rd position syllables (the 1st-3rd RT difference is greatest). Graded RT effect is not linear, and therefore cannot be ascribed simply to faster RTs in later stream positions. Black dots are estimated marginal means from linear model with block and pairings as predictors. Vertical lines represent 95% CIs. B. Mean RT differences between pairs of positions did not vary significantly by block, suggesting the general RT effect did not vary much over longer exposures. Center dots are estimated marginal means from the linear model, vertical lines represent 95% CIs. C. Exp. 1: RTs to targets in each stream position (averaged over ordinal positions and blocks) did not show a consistent downward trend. Dots represent mean RTs to targets in each stream position, red line indicates linear prediction, grey ribbon indicates SEM. D. Exp. 2: RTs to targets in each stream position for each condition (averaged over ordinal positions and trials) showed a quadratically-*

*shaped trend that increased in overall RT until target number 12-14, and thereafter began to decrease. This pattern is unlikely to have generated the graded RT effect we observed in our main RT by ordinal position analysis. Dots represent mean RTs to targets in each stream position, red line indicates linear prediction, grey ribbon indicates SEM. Solid line denotes structured condition, Dashed line denotes random condition. E. Stream position (1-216) of each target syllable in Experiment 1. Each target syllable appeared in virtually all possible positions in the stream. F. Median RT (in ms) for target syllables in Experiment 1 varied by triplet position (word-initial, -medial, or –final), but was independent of where in the stream (positions 1-216) the target appeared.*

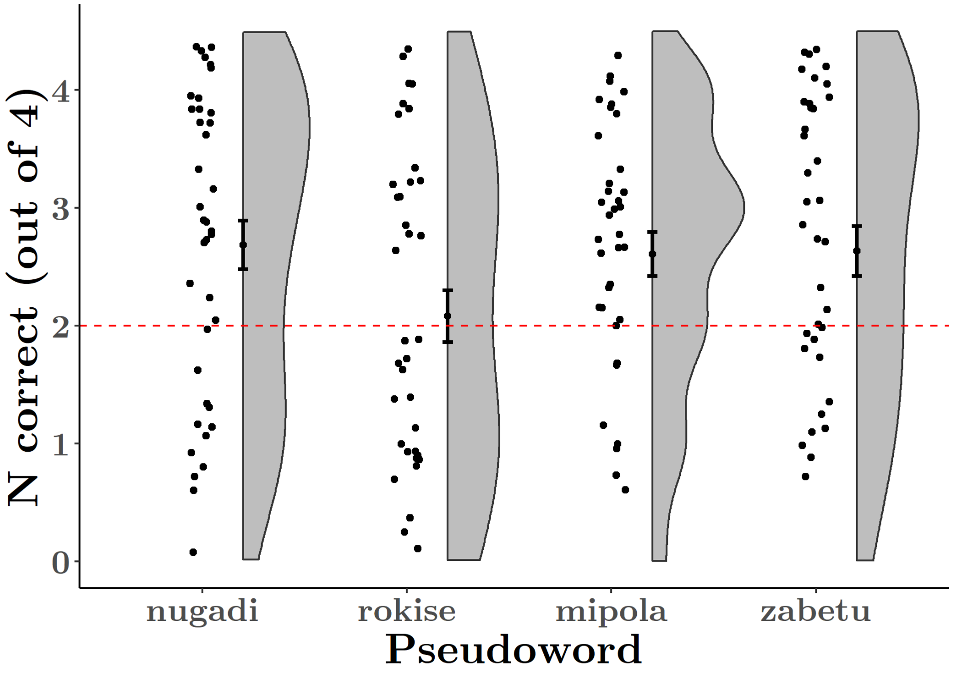

***Figure S3. Exp. 1: Preference for pseudowords over part-word foils was above chance for 3 of 4 pseudowords.*** *Number correct trials (out of 4 for each pseudoword) was above chance (2) for three of the four pseudowords. Jittered dots represent number correct trials for each individual. Adjacent black dot and error bars represent group mean and SEM. Overall word recognition performance was therefore not driven by successful discrimination of merely one word.*

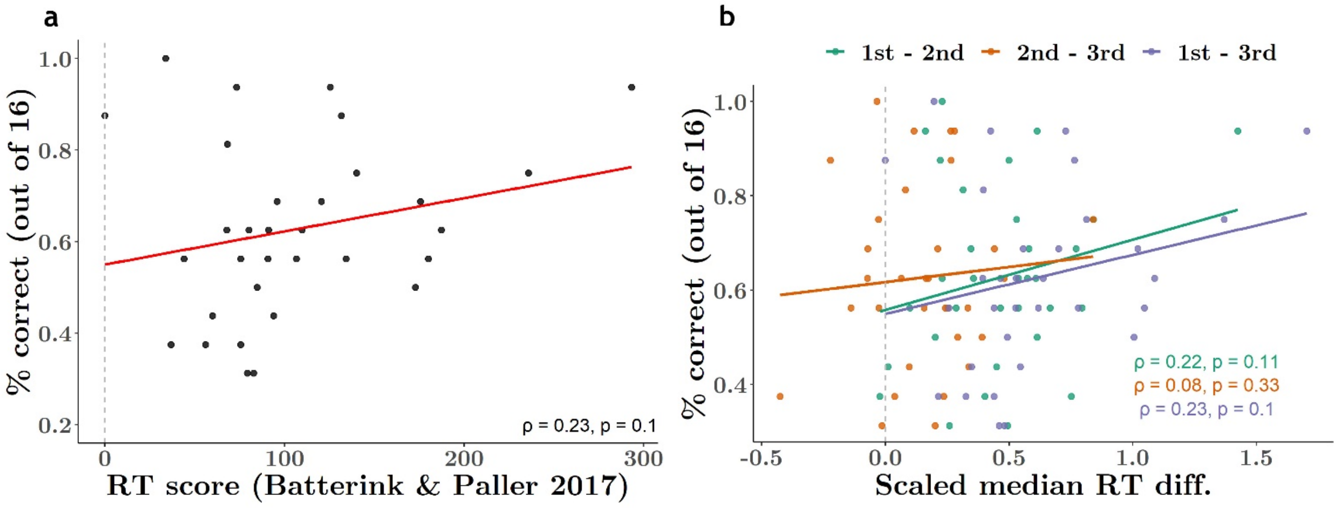

***Figure S4. Exp. 1: Correlation between online and offline SL measures following two exploratory methods reveals no significant relationship.*** *A. Word recognition (% correct) was correlated for each individual with an RT score, as per Batterink & Paller 2017, calculated by simple subtraction of median 1st-median 3rd position RT (Pearson's product-moment correlation). We found a weak correlation ($\rho = 0.23, p = 0.1$). B. Word recognition was correlated with three measures of online SL for each participant: we computed the difference in scaled median RTs between ordinal position separately. Scaling median RT values allowed all participants' scores to be more comparable. Calculating median differences for each pair allowed us to account for the possibility that e.g. the difference between 1st and 2nd positions better predicted word recognition performance than the difference between 2nd and 3rd position RTs. Median RT differences between all three pairs had only a weak relationship with word recognition performance. The strongest relationship existed between word recognition accuracy and 1st-3rd median RT ($\rho = 0.23, p = 0.1$), which revealed the exact same relationship as Batterink & Paller's method. Dashed grey line in both graphs represents threshold at which there was no difference in RTs between the respective position pair.*

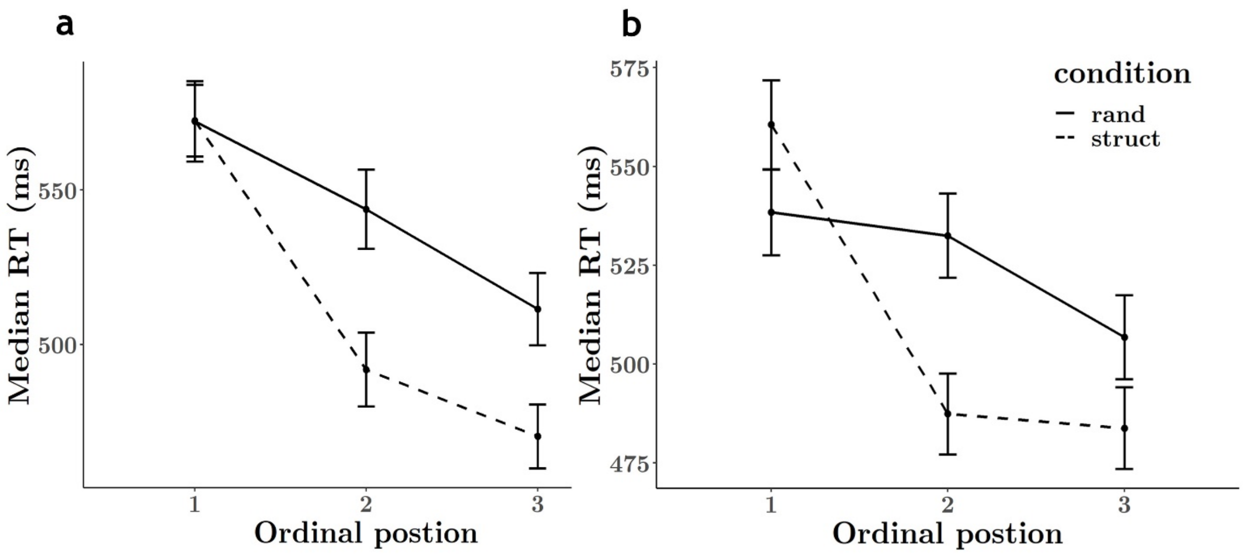

***Figure S5. Exp. 2: Graded RT effect for random and structured conditions in both between-subject condition orders.*** *A. Overall median RTs to targets in each ordinal position for participants who performed the structured condition first and the random condition second. B. Overall median RTs for participants who performed the random condition first and the structured condition second. Error bars represent 95% CIs. RT is strongly modulated by ordinal position in the structured condition in both between-subject orders. Those who performed the random condition first showed somewhat smaller differences in their RTs to each position (B, solid line) than those who performed this task after exposure to the structured stream (A, solid line). Despite observing faster responses to later ordinal positions in the random condition, the overall RT effect can be safely ascribed to the embedded regularities, as the RT effect is significantly larger in the structured condition (A-B, dashed line).*

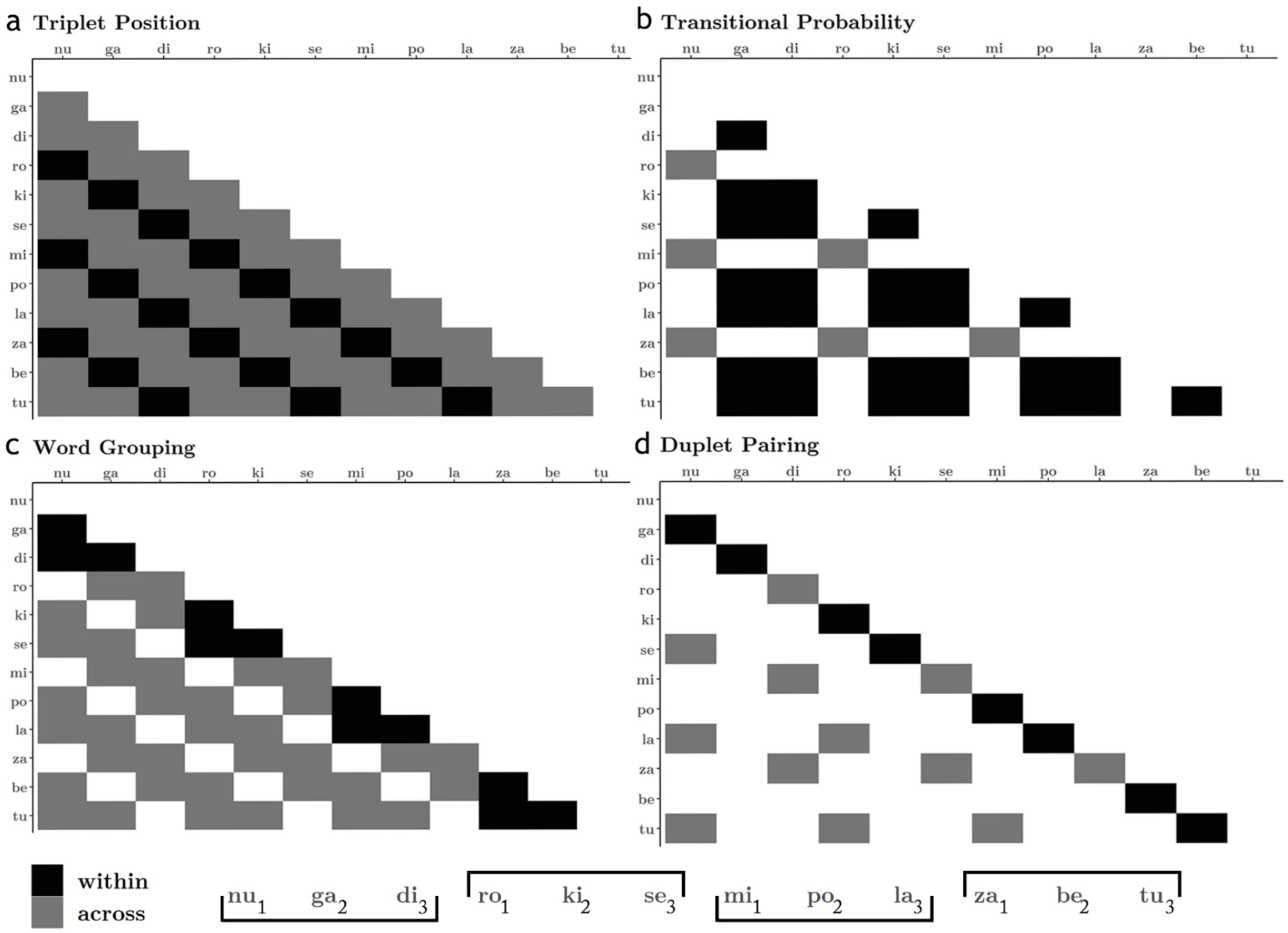

***Figure S6. Graphical representation of within and across group composition for each RSA analysis.*** Within *and* across *groups were created for each feature by selecting syllable pairs where the relationship between the two syllables either respected or violated the feature, respectively, or in the case of transitional probability, where both syllables either have a transitional probability of 1 (within) or of 0.33 (across). A. For the test of triplet position, within values were the correlations between syllables that occupy the same position in the pseudoword, and across values were correlations between syllable pairs of different triplet positions. B. For the test of transitional probability, within values were drawn from pairs of syllables representing the "inner part" of pseudowords (each having a TP of 0.33), and across values were drawn from pairs representing word boundaries (each having a TP of 1). C. For the test of word grouping, within values were drawn from all pairings of syllables where both syllables are members of the same pseudoword, and across values were from all possible "phantom word" combinations. D. For the test of duplet pairing, within values were drawn from all syllable pairings that constitute a part of a pseudoword, and across values were pairings of syllables where each syllable comes from a different pseudoword, while the triplet order of the syllables is respected. E.g. ga-se is an "across duplet" but ga-ki is not.*

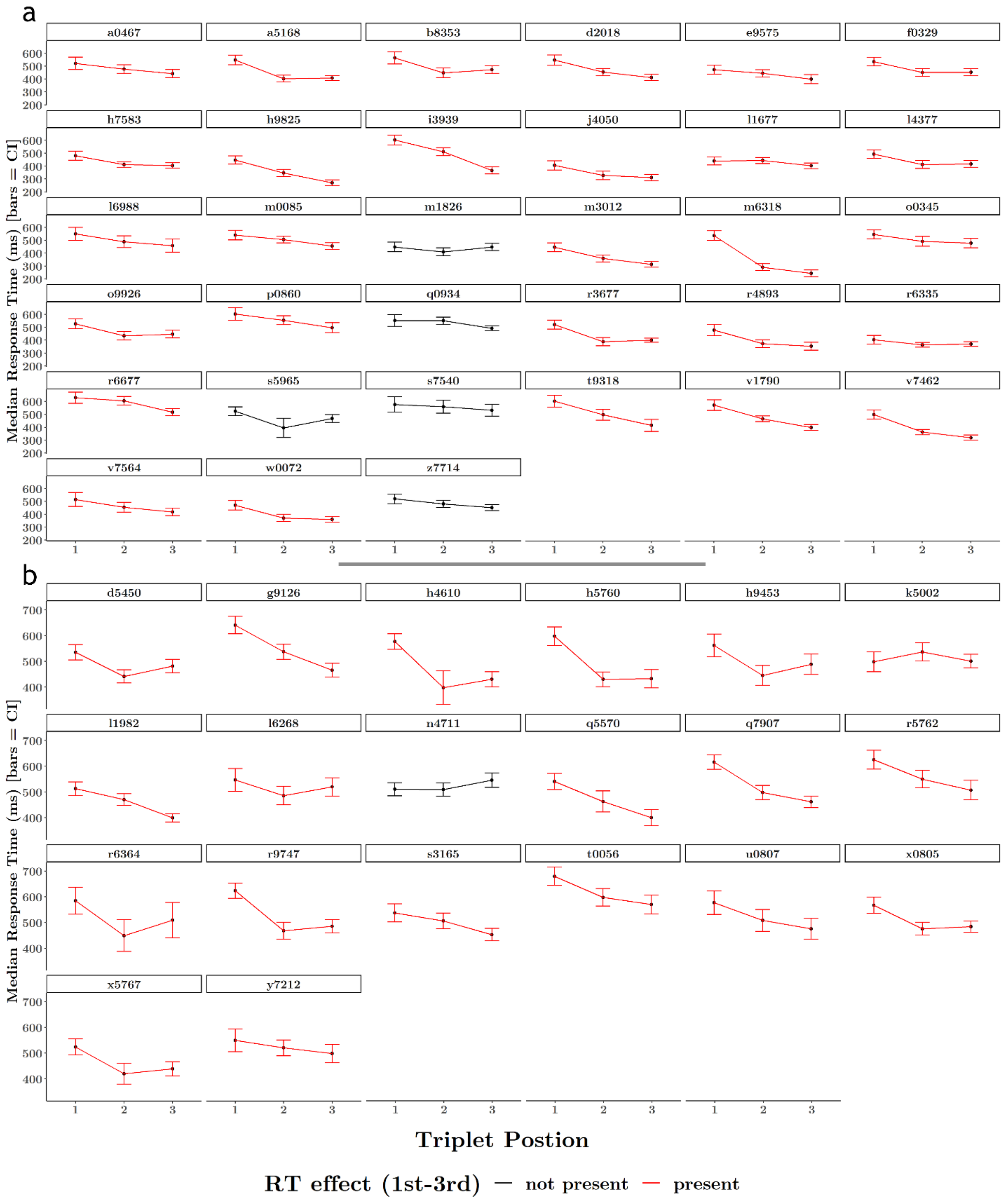

***Figure S7. Graded RT effect observable in most participants.*** *The graded RT effect is observable in 89% of participants. A participant was considered to display facilitated RTs in response to the statistical regularities of the stream if the mean difference between either position 1 and 2 syllables, or the mean difference between position 1 and 3 syllables, was significant at a 95% alpha level.* ***A.*** *Experiment 1. 28 of 33 participants exhibit the effect.* ***B****. Experiment 2. 19 of 20 participants exhibit the RT effect.*

### Supplementary Tables

**Table S1. GLM Results Experiment 1**

| | *Dependent variable:* | | |
|---|---|---|---|
| | reaction time (s) | | |
| | lesser | lesser (random slopes) | fuller |
| | (1) | (2) | (3) |
| Intercept(Pos 1) | -0.63*** (-0.68, -0.57) | -0.64*** (-0.68, -0.61) | -0.69*** (-0.76, -0.62) |
| Pos 2 | -0.16*** (-0.18, -0.14) | -0.14*** (-0.19, -0.10) | -0.11*** (-0.17, -0.05) |
| Pos 3 | -0.23*** (-0.25, -0.21) | -0.22*** (-0.28, -0.15) | -0.18*** (-0.24, -0.12) |
| Block 2 | | | 0.07** (0.01, 0.13) |
| Block 3 | | | 0.08*** (0.02, 0.15) |
| Block 4 | | | 0.08** (0.02, 0.14) |
| Block 5 | | | 0.11*** (0.05, 0.17) |
| Block 6 | | | 0.03 (-0.03, 0.09) |
| Block 7 | | | 0.05 (-0.01, 0.11) |
| Block 8 | | | 0.06* (-0.005, 0.12) |
| Pos 2:Block 2 | | | -0.04 (-0.12, 0.04) |
| Pos 3:Block 2 | | | -0.03 (-0.11, 0.05) |
| Pos 2:Block 3 | | | -0.07* (-0.16, 0.01) |
| Pos 3:Block 3 | | | -0.03 (-0.11, 0.05) |
| Pos 2:Block 4 | | | -0.05 (-0.13, 0.04) |
| Pos 3:Block 4 | | | -0.07 (-0.15, 0.01) |
| Pos 2:Block 5 | | | -0.06 (-0.14, 0.03) |
| Pos 3:Block 5 | | | -0.14*** (-0.22, -0.06) |
| Pos 2:Block 6 | | | -0.01 (-0.10, 0.07) |
| Pos 3:Block 6 | | | 0.01 (-0.06, 0.09) |
| Pos 2:Block 7 | | | -0.06 (-0.15, 0.02) |
| Pos 3:Block 7 | | | -0.09** (-0.17, -0.004) |
| Pos 2:Block 8 | | | -0.06 (-0.14, 0.03) |
| Pos 3:Block 8 | | | -0.06 (-0.14, 0.02) |
| Fixed Effects | Subject | Position \| Subject | Subject |
| Fixed Effects Struct. | Rand. Int. | Rand. Int., Slope | Rand Int. |
| Observations | 9,531 | 9,531 | 9,531 |
| Log Likelihood | 3,199.61 | 3,312.13 | 3,224.15 |
| Akaike Inf. Crit. | -6,389.23 | -6,604.26 | -6,396.29 |
| Bayesian Inf. Crit. | -6,353.42 | -6,532.64 | -6,210.07 |
| *Note:* | | | *p**p***p<0.01 |
| | | | Fitted using Gamma distribution and log link function. |

**Table S2. GLM Results Experiment 2**

| | *Dependent variable:* |
|---|---|
| | reaction time (s) |
| Intercept(Pos 1/Rand) | -0.54$^{***}$ (-0.58, -0.50) |
| Struct | 0.002 (-0.04, 0.04) |
| Pos 2 | -0.03$^{**}$ (-0.05, -0.01) |
| Pos 3 | -0.08$^{***}$ (-0.10, -0.06) |
| Struct:Pos 2 | -0.11$^{***}$ (-0.14, -0.08) |
| Struct:Pos 3 | -0.09$^{***}$ (-0.12, -0.06) |
| Fixed Effects | Condition Order/Subject + Condition |
| Fixed Effects Struct. | Rand. Int. + Rand. Slope |
| Observations | 6,524 |
| Log Likelihood | 4,140.28 |
| Akaike Inf. Crit. | -8,250.56 |
| Bayesian Inf. Crit. | -8,148.81 |
| *Note:* | $^{*}$p$^{**}$p$^{***}$p<0.01 |
| | Fitted using Gamma distribution and log link function. |

**Table S3. Experiment 1 Detection Accuracy**

**A**

| word | N | mean | sd | se | ci |
|---|---|---|---|---|---|
| nugadi | 3,425 | 0.772 | 0.419 | 0.007 | 0.014 |
| rokise | 3,378 | 0.722 | 0.448 | 0.008 | 0.015 |
| mipola | 3,452 | 0.727 | 0.445 | 0.008 | 0.015 |
| zabetu | 3,306 | 0.586 | 0.493 | 0.009 | 0.017 |

**B**

| target | N | mean | sd | se | ci |
|---|---|---|---|---|---|
| be | 1,015 | 0.449 | 0.498 | 0.016 | 0.031 |
| di | 1,154 | 0.824 | 0.381 | 0.011 | 0.022 |
| ga | 1,149 | 0.727 | 0.446 | 0.013 | 0.026 |
| ki | 1,150 | 0.804 | 0.397 | 0.012 | 0.023 |
| la | 1,152 | 0.814 | 0.389 | 0.011 | 0.022 |
| mi | 1,136 | 0.583 | 0.493 | 0.015 | 0.029 |
| nu | 1,122 | 0.766 | 0.424 | 0.013 | 0.025 |
| po | 1,164 | 0.783 | 0.413 | 0.012 | 0.024 |
| ro | 1,082 | 0.486 | 0.500 | 0.015 | 0.030 |
| se | 1,146 | 0.861 | 0.346 | 0.010 | 0.020 |
| tu | 1,148 | 0.761 | 0.426 | 0.013 | 0.025 |
| za | 1,143 | 0.531 | 0.499 | 0.015 | 0.029 |

| Table S4. Experiment 2 Detection Accuracy | | | | | | |
|---|---|---|---|---|---|---|
| condition | target | N | mean | sd | se | ci |
| random | be | 316 | 0.737 | 0.441 | 0.025 | 0.049 |
| | di | 333 | 0.883 | 0.322 | 0.018 | 0.035 |
| | ga | 326 | 0.709 | 0.455 | 0.025 | 0.050 |
| | ki | 318 | 0.852 | 0.355 | 0.020 | 0.039 |
| | la | 314 | 0.828 | 0.378 | 0.021 | 0.042 |
| | mi | 313 | 0.744 | 0.437 | 0.025 | 0.049 |
| | nu | 322 | 0.807 | 0.395 | 0.022 | 0.043 |
| | po | 317 | 0.817 | 0.387 | 0.022 | 0.043 |
| | ro | 313 | 0.776 | 0.417 | 0.024 | 0.046 |
| | se | 349 | 0.880 | 0.326 | 0.017 | 0.034 |
| | tu | 324 | 0.722 | 0.449 | 0.025 | 0.049 |
| | za | 305 | 0.777 | 0.417 | 0.024 | 0.047 |
| structure | be | 349 | 0.682 | 0.466 | 0.025 | 0.049 |
| | di | 346 | 0.925 | 0.264 | 0.014 | 0.028 |
| | ga | 345 | 0.791 | 0.407 | 0.022 | 0.043 |
| | ki | 344 | 0.875 | 0.331 | 0.018 | 0.035 |
| | la | 342 | 0.827 | 0.378 | 0.020 | 0.040 |
| | mi | 343 | 0.872 | 0.335 | 0.018 | 0.036 |
| | nu | 333 | 0.877 | 0.329 | 0.018 | 0.035 |
| | po | 340 | 0.918 | 0.275 | 0.015 | 0.029 |
| | ro | 331 | 0.801 | 0.400 | 0.022 | 0.043 |
| | se | 347 | 0.827 | 0.379 | 0.020 | 0.040 |
| | tu | 340 | 0.862 | 0.346 | 0.019 | 0.037 |
| | za | 344 | 0.869 | 0.338 | 0.018 | 0.036 |
